# Janzen-Connell effects and habitat-induced aggregation synergistically promote species coexistence

**DOI:** 10.1101/2022.04.11.487940

**Authors:** Daniel J. B. Smith

## Abstract

Janzen–Connell (JC) effects, habitat partitioning (HP), and dispersal limitation are three processes thought to shape tropical forest diversity and patterns of species aggregation, yet how they jointly influence species coexistence remains poorly understood. I analyze a spatially explicit model that incorporates all three processes, tracking propagules subject to habitat filtering and conspecific density dependence. The model shows that dispersal limitation and habitat filtering interact with JC-effects in qualitatively different ways. Aggregation driven by dispersal limitation weakens the ability of JC-effects to maintain coexistence, whereas when JC-effects and HP operate together in spatially autocorrelated habitats, richness increases with conspecific aggregation. JC-HP interactions strongly depend on habitat spatial structure. If habitats are not spatially autocorrelated, JC-effects and HP minimally interact (or act antagonistically when coupled with dispersal limitation) and do not maintain high species richness. However, when habitats are positively autocorrelated the two mechanisms act synergistically. To explain these patterns, I introduce a novel metric — the spatial JC–HP covariance — which quantifies how strongly JC-effects are concentrated in favorable versus unfavorable environments, providing a direct measure of the extent to which JC-effects and HP synergistically promote coexistence.

## Introduction

Explaining how tropical forests sustain extraordinary levels of species diversity remains a central challenge in ecology. The difficulty stems from the competitive exclusion principle: without interspecific niche differences, the species with the highest fitness should exclude all others (Hardin, 1960; Levine & HilleRisLambers, 2009). This puzzle has motivated decades of empirical and theoretical work to identify the mechanisms that allow stable coexistence.

Two spatial coexistence mechanisms —– Janzen–Connell (JC) effects and habitat partitioning (HP) –— are frequently invoked to explain high tree diversity in tropical forests (Fig. 1A). JC-effects occur when host-specific enemies reduce juvenile survival near conspecific adults, causing survival to decline with increasing conspecific density. This density-dependent mortality generates negative frequency dependence: rare species gain a per-capita advantage because their enemies are also rare. JC-effects have been documented in forests worldwide (e.g. Hyatt *et al*., 2003; Mangan *et al*., 2010; Comita *et al*., 2014; Song *et al*., 2021; Hülsmann *et al*., 2024). By contrast, habitat partitioning occurs when species differ in their responses to the abiotic environment (e.g. topography, soil), such that each species performs best in a subset of habitats (Chesson, 2000a; Snyder & Chesson, 2003; Stein *et al*., 2014). Many studies — typically measuring the effects of conspecific density and habitat conditions on seedling and sapling survival — find both processes occur simultaneously (e.g. Uriarte *et al*., 2004; Freckleton & Lewis, 2006; Queenborough *et al*., 2009; Bai *et al*., 2012; Piao *et al*., 2013; Lu *et al*., 2015; Wu *et al*., 2016; Johnson *et al*., 2017; Pu *et al*., 2017; Cao *et al*., 2018; Yao *et al*., 2020; Magee *et al*., 2021; Huang *et al*., 2022). It is therefore important to elucidate how they interact.

**Figure 1:**
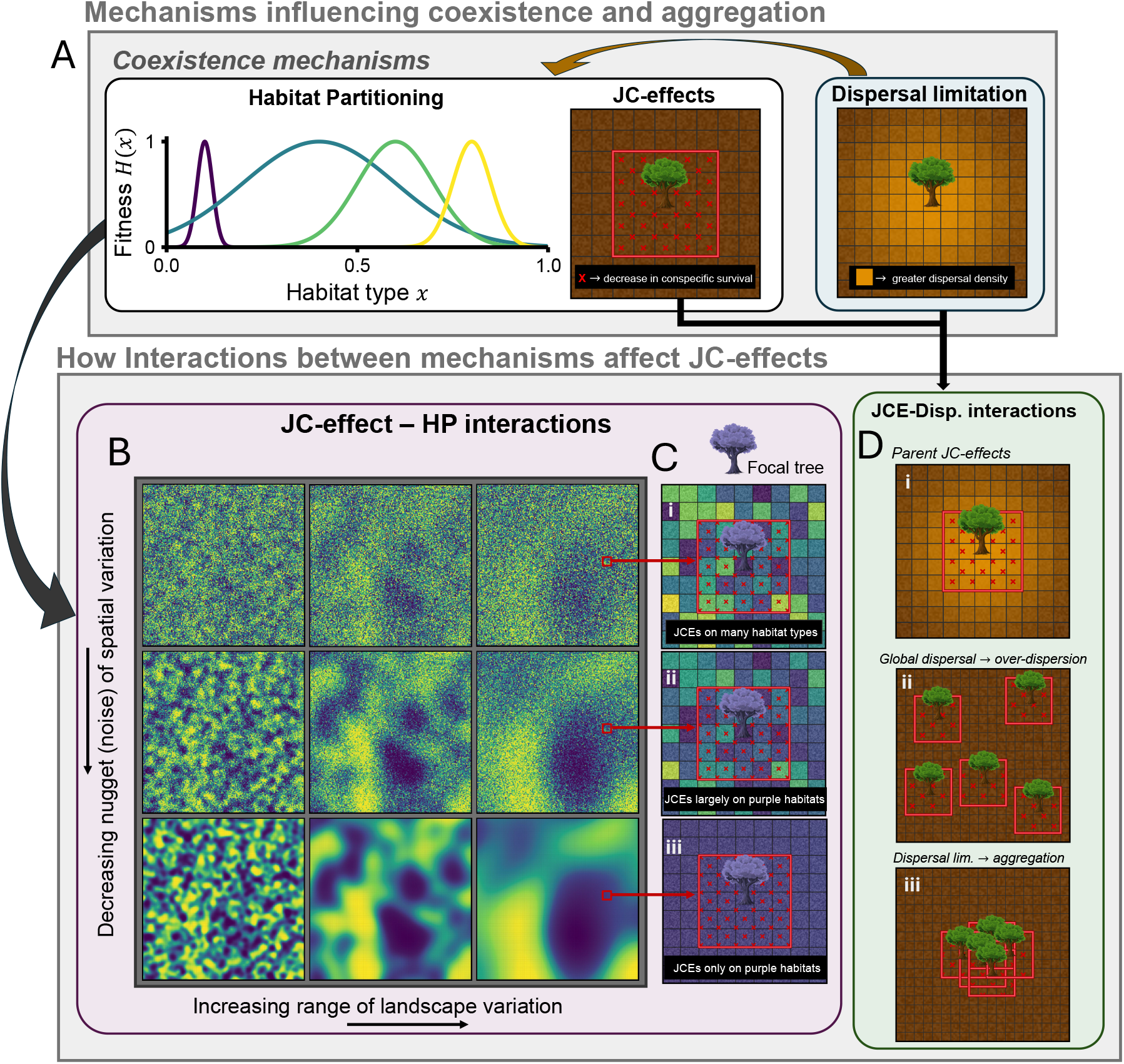
Conceptual overview of coexistence mechanisms and their interactions. (A) “Habitat partitioning” shows species responses to the environment *H*(*x*) (*σ*_*h*_ = 0.02, 0.2, 0.1, and 0.05, left to right). “JC-effects” depict the *M* × *M* Moore neighborhood of influence (red) around a focal tree, *M* = 5. “Dispersal limitation” shows the relative propagule pressure from a focal tree (orange shading). (B) Spatial distributions of habitat under varying nugget and range parameters. The nugget controls fine-scale heterogeneity (i.e., an uncorrelated microscale white-noise variance term added at each location); the range controls the size of habitat patches. Nugget values are 10, 1, and 0 (top to bottom) and range values are 0.05, 0.2, 0.5 (left to right). (C) depicts JC–HP interactions: for a focal tree positively associated with purple habitat patches, the nugget affects which patch types it induces JC-effects on (i-iii). (D) JC-effects interact with dispersal. (i) Locally dispersed propagules are subject to its parent’s JC-effects. (ii) Under global dispersal, JC-effects generate overdispersion. (iii) With dispersal limitation, conspecifics may aggregate.

Although both coexistence mechanisms ultimately maintain diversity by concentrating intraspecific competition relative to interspecific competition (Chesson, 2000b), they shape spatial distributions in opposite ways. Habitat partitioning aggregates species into favorable environments (e.g. Andersen *et al*., 2010; Jones *et al*., 2011; Brown *et al*., 2013; John *et al*., 2007; Potts *et al*., 2002; Umarani *et al*., 2024). In contrast, JC-effects generate conspecific repulsion and overdispersion of adults (Fig. 1D, ii; Chave *et al*., 2002; Wiegand & Moloney, 2013; Kalyuzhny *et al*., 2023). Because these mechanisms push spatial patterns in opposite directions, their joint operation may yield non-additive interactions that affect species coexistence in counter-intuitive ways.

Empirical studies account for this tension. Tree species are almost universally aggregated in tropical forests (Condit *et al*., 2000), a pattern largely (although not exclusively) driven by habitat specialization and dispersal limitation (Baldeck *et al*., 2013). These aggregation-generating processes complicate empirical detection of negative density dependence: the positive effects of favorable habitats can mask or counteract mortality generated from JC-effects (Wright, 2002; Chen *et al*., 2010; Bagchi *et al*., 2011; Wu *et al*., 2016; Liu *et al*., 2022; Feng *et al*., 2023). Dispersal limitation adds further aggregation that is independent of habitat heterogeneity. As a result, spatial point-pattern analyses must disentangle overlapping influences of dispersal, habitat filtering, and density-dependent mortality (Detto & Muller-Landau, 2013; Zhu *et al*., 2013). Recent analyses at the Barro Colorado Island (BCI) forest dynamics plot illustrate this challenge: strong conspecific repulsion becomes evident only after controlling for aggregation caused by dispersal limitation (Kalyuzhny *et al*., 2023).

Despite empirical advances in isolating the signal of negative density dependence from dispersal limitation and habitat specialization, theory has not clarified how their joint operation —– and the aggregation patterns they produce — shape species coexistence. Conspecific aggregation can either promote or hinder species richness, depending on the context (Chesson & Neuhauser, 2002; Murrell *et al*., 2002; Bolker *et al*., 2003). For example, under density dependence, coexistence is hindered when rare species are more intraspecifically aggregated than common species (Stump & Comita, 2020), but enhanced when rare species are less intraspecifically aggregated than common species (Wiegand *et al*., 2025). Aggregation driven by dispersal also often positively influences the ability of habitat partitioning to maintain coexistence (Snyder, 2008; D’Andrea & O’Dwyer, 2021). Although both dispersal limitation and habitat specialization generate aggregation, they do so through distinct pathways: dispersal limitation produces clustering largely independent of habitat heterogeneity, whereas habitat specialization concentrates individuals in locations where they experience high fitness. Because JC-effects modify local environments, these contrasting forms of aggregation may interact with JC-effects in different ways (at least when habitats are positively spatially autocorrelated; Fig. 1C vs. Fig. 1D, sub-panel iii). Yet this possibility has received little theoretical attention. The key question is: How do these different forms of aggregation interact with the repulsion generated by JC-effects to shape species coexistence?

Ultimately, the crucial issue is whether interactions between JC-effects and habitat partitioning act synergistically or antagonistically — especially given that past work casts doubt on whether either mechanism alone can explain the extraordinary diversity of tropical forests. Although previous theory suggests that JC-effects alone can maintain high diversity in otherwise neutral communities (Levi *et al*., 2019), JC-effects are likely insufficient when species differ in fitness. For example, Chisholm & Fung (2020) argue that empirically estimated fitness differences can substantially erode species richness, using interspecific variation in fecundity and low-density seedling recruitment rates from the BCI forest dynamics plot as proxies for fitness. Importantly, however, these proxies may overstate true fitness differences if seed production trades off against other life-history components (e.g., with survival and stress tolerance; Muller-Landau, 2010). Nevertheless, coexistence mechanisms must be sufficiently strong to offset fitness differences (Chesson, 2000b); the extent to which JC-effects meet this requirement remains contentious.

A parallel concern applies to habitat partitioning: although it shapes species’ spatial distributions and demography, the low dimensionality of abiotic variation relative to the number of species in tropical forests makes habitat partitioning alone unlikely to explain observed diversity (Wright, 2002; Silvertown, 2004; Terborgh, 2015). The limitations of each mechanism in isolation raise a central question: Do JC-effects and habitat partitioning, acting together, maintain more or less diversity than expected from the sum of their individual contributions? And how does dispersal limitation modify the outcome of this interaction?

Synergistic interactions would provide a novel explanation for the maintenance of coexistence, whereas antagonistic interactions would deepen the mystery of what sustains diversity. These possibilities likely hinge on the spatial structure of populations. Because seedling (or sapling) survival under JC-effects is a non-linear function of local conspecific density (e.g. Detto *et al*., 2019), the contribution of JC-effects to coexistence depends on how individuals are distributed in space. In particular, aggregation increases variance in conspecific neighborhood density; with non-linear density dependence, this can shift the spatial average of mortality caused by JC-effects, even if mean density is fixed (i.e. Jensen’s inequality; Jensen, 1906). Therefore, the extent to which habitat specialization and dispersal limitation generate aggregation may amplify or dampen both the local strength of density dependence (Chen *et al*., 2010; Fortunel *et al*., 2018) and its downstream community-wide consequences.

I analyze a spatially explicit model incorporating JC-effects, habitat partitioning, and dispersal limitation, while allowing species to differ in intrinsic fitness (e.g., via fecundity and other density-independent demographic components). I evaluate the ability of JC-effects and habitat partitioning, alone and in combination, to maintain species richness, with particular attention to how aggregation arising from dispersal limitation versus habitat specialization modifies the contribution of JC-effects to coexistence. I examine several scenarios, including a stress–tolerance–fecundity trade-off (in which seed size mediates a trade-off between fecundity and juvenile survival (tolerance) across spatially heterogeneous habitat conditions; Muller-Landau, 2010) and interspecific variation in susceptibility to JC-effects. Across scenarios, dispersal limitation weakens the ability of JC-effects to maintain diversity. In contrast, habitat-driven aggregation acts synergistically with JC-effects to promote coexistence, often exceeding what would be expected from the sum of their individual effects. This synergy is captured by the spatial covariance between JC-effects and habitat partitioning (the “JC–HP covariance”): species richness is highest when trees disproportionately generate JC-effects in the habitat types where they also perform best. Because this covariance can be estimated from spatial demographic data, it provides a direct link between tree spatial patterns and the processes that maintain tropical forest diversity.

## Methods

### Model description

I use a discrete-time site occupancy model similar to those developed by Levi *et al*. (2019), Chisholm & Fung (2020), and Smith (2022). The model tracks the dynamics of a local forest community composed of *P* patches, each occupied by a single adult tree. The local community contains *N* species, which draw from a regional pool of *L* species (*L* ≥ *N*). Each time step proceeds through four stages: (1) Adults disperse propagules (i.e., seeds) throughout the landscape. (2) Propagules experience mortality due to habitat effects and JC-effects. (3) Each adult dies with probability *δ*. (4) Vacant patches are colonized either by a propagule from the local seed rain or by an immigrant drawn from the regional species pool. See Supplementary Table 1 for a list of all parameters and variables.

To model propagule arrival, let *x* denote a patch (i.e., a discrete grid cell) in space. At each time step, species *i*(*i* = 1, 2, …, *N*) disperses propagules across the landscape in proportion to its intrinsic fitness parameter *Y*_*i*_. This parameter sets baseline propagule input (and hence baseline recruitment weights) and thus captures density-independent fitness differences among species as a composite of demographic components (e.g., fecundity and baseline establishment/survival), prior to habitat filtering and JC-induced mortality. Dispersal occurs either globally, with propagules distributed uniformly across all patches, or locally according to a Gaussian kernel with standard deviation *σ*_*D*_ (in grid-cell units), which determines the spatial scale of dispersal. The expected propagule density of species *i* arriving at patch *x* is denoted *d*_*i*_(*x*). Full details of the dispersal kernel are provided in the Supplementary Information (see “General model derivation and details”).

After propagules arrive, their survival depends on two factors: abiotic habitat effects and JC-effects. I begin by describing habitat effects. Each patch *x* is assigned a habitat value *E*(*x*) ∈ [0, 1] that influences species’ demographic performance. The survival of species *i*’s seedlings on patch *x* is given by a Gaussian response function of the local habitat value:

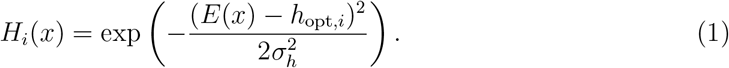

Here, *h*_opt,*i*_ is species *i*’s habitat optimum (0 ≤ *h*_opt,*i*_ ≤ 1), and *σ*_*h*_ determines niche breadth; smaller *σ*_*h*_ indicates greater specialization. The function *H*_*i*_(*x*) ranges between 0 and 1 and captures the relative survival probability of propagules across the habitat axis.

In the main text, I examine the above Gaussian habitat response, in which all species share the same value of *σ*_*h*_. However, species may also display specialist–generalist trade-offs and exhibit non-Gaussian environmental responses. To test if results were robust to this possibility, I also performed analyses based on the so-called “stress–tolerance–fecundity trade-off” (Muller-Landau, 2010; D’Andrea *et al*., 2013; D’Andrea & O’Dwyer, 2021), in which seed size mediates a trade-off between fecundity and tolerance of stressful environments (e.g., drought or low resource availability; Moles *et al*., 2003; Moles & Westoby, 2004). The stress–tolerance–fecundity trade-off model and related results are described in the Supplementary Information (“Tolerance–fecundity trade-off model”).

Propagule survival is further reduced by JC-effects, representing mortality caused by nearby conspecific adults (e.g., via host-specific natural enemies). JC-effects act within an *M* × *M* Moore neighborhood around each patch *x* (including diagonals; Fig. 1A). Let *A*_*i*_(*M*(*x*)) denote the number of conspecific adults of species *i* within this neighborhood. The probability that propagules of species *i* on patch *x* die due to JC-effects is

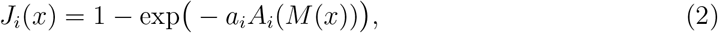

where *a*_*i*_ is the baseline strength of JC-effects. I do not model natural enemies explicitly; instead, Eq. (2) assumes exponential survival in which conspecific adults contribute additively to the cumulative hazard (i.e., to − log(survival)), with each conspecific adult in *M*(*x*) adding hazard *a*_*i*_ to the focal patch *x*. I refer to the spatially summed hazard contributed by a single adult across its *M* × *M* footprint as the “JC-effect pressure”, equal to *a*_*i*_*M*^2^ (see Supplementary Information “General model derivation and details”).

Combining dispersal, habitat effects, and JC-effects, the expected number of viable propagules of species *i* on patch *x* is

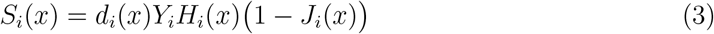

where survival after JC-effects is represented by 1 − *J*_*i*_(*x*). Thus, a species’ recruitment on a patch depends on its dispersal pressure (*d*_*i*_(*x*)), its intrinsic fitness *Y*_*i*_, habitat suitability *H*_*i*_(*x*), and its probability of surviving JC-effects 1 − *J*_*i*_(*x*).

Following propagule dispersal and survival, adult dynamics unfold. At each time step, every adult dies with probability *δ*. Vacant patches are then filled via a lottery, in which the replacement individual is drawn either from propagules produced within the modeled landscape of *P* patches (local recruitment) or, with small probability, from an external regional species pool (immigration). With probability 1 − *ν*, recruitment is local and species *i* claims patch *x* with probability

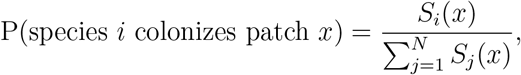

where *S*_*i*_(*x*) is the number of surviving propagules from Equation (3) (which incorporates dispersal within the modeled landscape). With probability *ν*, the recruit is instead drawn from the regional species pool, representing rare long-distance dispersal/background seed rain that allows locally absent species to re-enter. When immigration occurs, establishment on patch *x* is still filtered through the same habitat and JC-effect survival terms (so, conditional on immigration, the establishing species at patch *x* is still biased by habitat suitability and local JC pressure at *x*). Introducing *ν >* 0 yields a well-defined dynamic equilibrium (preventing the eventual absorbing state of monodominance when *ν* = 0).

### Parameterizations for simulations

I used empirical data — largely from the 50-ha Forest Dynamics Plot at BCI — to roughly parameterize key quantities within a biologically feasible range.

In the model, *M* is the side length (in patches) of the Moore neighborhood within which conspecific adults reduce offspring survival, and it is set by (i) the biological interaction radius *R* (meters, m) and (ii) the density of reproductive adults, *g* (per meters squared, m^−2^). Field studies suggest JC-effects often decline beyond ∼10–20 m (Hubbell *et al*., 2001; Johnson *et al*., 2014), though they can extend further (Kalyuzhny *et al*., 2023); here I assume a more modest *R* ∼ 5–15 m. At BCI, published counts of reproductive adults (∼30,000– 50,000 across 50 ha, depending on maturity cutoffs) imply *g* ∼ 0.06–0.10 m^−2^ (Comita *et al*., 2007; D’Andrea & O’Dwyer, 2021). In the square-lattice geometry, each patch has area 1*/g* and side length 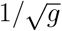, so a neighborhood of width *M* spans a 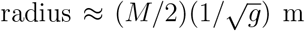, giving 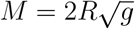. Using the ranges above yields *M* ≈ 3–9. Accordingly, (in all but one set of simulations) I assumed *M* ∈ {1, 3, 5, 7, 9}, spanning small to relatively large neighborhood effects.

I parameterize the baseline strength of JC-effects, *a*, using empirical estimates of neighborhood mortality. Meta-analyses suggest that seedling survival near a conspecific adult is reduced by roughly 60%, which corresponds to *a* ≈ 1 under the exponential mortality function (since a single nearby conspecific implies mortality 1 − *e*^−*a*^, and 1 − *e*^−1^ ≈ 0.63) (Comita *et al*., 2014; Song *et al*., 2021). As a default for most simulations, I use a conservative value of *a* = 0.5 (implying 1 − *e*^−0.5^ ≈ 0.39 mortality when *A*_*i*_ = 1), so that any coexistence generated by JC–HP interactions does not rely on exceptionally strong neighborhood mortality. This assumption is particularly conservative because JC-effects in the model aggregate mortality from seed arrival through establishment, whereas the estimates above are typically based on seedling survival. I therefore also examined larger values of *a* in sensitivity analyses (see below). Unless otherwise noted, species were assumed to share the same baseline JC-effect strength. Because empirical studies document substantial interspecific variation in sensitivity to conspecific density (Comita *et al*., 2010; Song *et al*., 2021), I additionally ran simulations in which *a* varied among species, with values drawn from a Gamma distribution (see Supplementary Information “Interspecific variation in baseline JC-effect strength”).

Substantial interspecific variation in fitness-related demographic rates (e.g., fecundity and density-independent survival) is well documented in tropical forests such as BCI. In particular, fecundity and related rates are highly right-skewed and vary widely among species (Dalling *et al*., 2002; Wright *et al*., 2005; Chisholm & Fung, 2020). Following Chisholm & Fung (2020) and D’Andrea & O’Dwyer (2021), I represent intrinsic fitness (*Y*) as a log-normally distributed variable, *Y* ∼ lognormal(*µ* = 0, *σ*_*Y*_). A lognormal form is natural here because *Y* is strictly positive and can be interpreted as a multiplicative composite of underlying density-independent demographic components (e.g., fecundity × baseline establishment/survival × low-density recruitment probability), which yields right-skewed variation among species. For most simulations, I adopt *σ*_*Y*_ = 0.3, which produces roughly an order of magnitude of variation in fitness among species — sufficient to test the robustness of coexistence mechanisms.

Species’ responses to abiotic heterogeneity — quantified by the niche-breadth parameter *σ*_*h*_ — are difficult to estimate directly because they integrate over multiple environmental factors that are rarely directly measured for most tropical tree species. I therefore explored a broad range of *σ*_*h*_ values in simulations, spanning from *σ*_*h*_ = 0.02 to *σ*_*h*_ = 0.2. Lower values (e.g., *σ*_*h*_ = 0.02; Fig. 1A, blue curve) represent relatively specialized species, while higher values (e.g., *σ*_*h*_ = 0.2; Fig. 1A, red curve) represent generalized responses. In simulations, each species’ habitat optimum, *h*_opt,*i*_, was drawn uniformly on [0, 1]. As noted earlier, I also considered an alternative formulation in which species experience a stress–tolerance–fecundity trade-off to test robustness (Supplementary Information).

I explored a range of dispersal limitation values, characterized by the dispersal kernel standard deviation *σ*_*D*_, from highly limited local dispersal (*σ*_*D*_ = 1) to global dispersal (*σ*_*D*_ = ∞). Throughout simulations, I explored *σ*_*D*_ ∈ {1.0, 1.5, 4.5, 6.25, 8.0, ∞}. All species were assumed to share the same value of *σ*_*D*_.

I set immigration to a low value, *ν* = 3 × 10^−6^, chosen so that an otherwise neutral baseline (no JC-effects, no habitat partitioning, and no fitness differences) supports only ∼ 1– 3 species at equilibrium (modal richness = 1). Richness substantially above this baseline therefore reflects the effects of JC-effects, habitat partitioning, or their interaction rather than immigration alone.

Simulations were run on a 175 × 175 grid, yielding ∼ 30,000 patches, each occupied by a single adult tree — comparable to the number of reproductive adults at BCI. Each patch was assigned a habitat value between 0 and 1. To investigate the effects of habitat-driven aggregation, I generated spatial habitat landscapes as Gaussian random fields using the Julia package GaussianRandomFields (Robbe, 2023), and rescaled values to a uniform [0, 1] distribution (preserving spatial ranks). These fields are parameterized by three quantities: the sill, the range, and the nugget. The sill sets the total variance of the field at large distances (fixed at 1 here for convenience), while the range specifies the distance at which sites become effectively uncorrelated (Fig. 1B, left to right) — formally, the range is the point where the variogram rises to 95% of the sill (Cressie, 1993; D’Andrea & O’Dwyer, 2021). With the parameterizations used here, a range of e.g. 0.5 implies the variogram reaches 95% of the sill at a distance of 50% of the total landscape side length. The nugget adds microscale white-noise variance to each point, producing local roughness without altering large-scale patchiness (Fig. 1B, top to bottom). Overall, the range controls the size of habitat patches, while the nugget controls their fine-scale heterogeneity.

### Aggregation metrics

In several parts of the analysis, I quantify species aggregation using two complementary metrics. First, as a simple measure of intraspecific spatial aggregation of adult trees in simulations, I computed 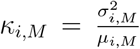 where *µ*_*i,M*_ and 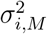 are the mean and variance of species *i*’s abundance across all *M* × *M* Moore neighborhoods. Here *µ*_*i,M*_ = *p*_*i*_*M*^2^, with *p*_*i*_ the proportion of patches occupied by species *i*. This local variance–to–mean ratio satisfies *κ*_*i,M*_ = 1 under spatial randomness, *κ*_*i,M*_ *>* 1 under aggregation, and *κ*_*i,M*_ *<* 1 under regular spacing. As a community-level summary, I use the average

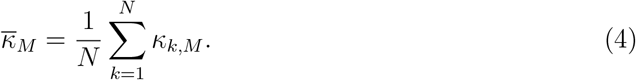

Because the scale *M* matches the neighborhood scale of JC-effects, this metric quantifies the outcome of interactions between the aggregation-generating processes (habitat partitioning and dispersal limitation) and the local repulsion imposed by JC-effects.

Second, I consider the pair-correlation function, which is the scale–specific counterpart of Ripley’s *K*–function (Ripley, 1976) and commonly used in spatial point pattern analyses (Wiegand & Moloney, 2013). For a point process of constant intensity *ψ*, the *K*-function is defined so that, for a typical individual (of a focal species), the expected number of other individuals within distance *r* is *ψK*(*r*). Under complete spatial randomness (CSR; individuals are located independently and uniformly at random in space), *K*(*r*) = *πr*^2^, while values above or below this benchmark indicate aggregation or inhibition, respectively. The pair-correlation function *g*(*r*) is obtained from *K*(*r*) as 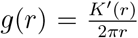, so that *g*(*r*) measures the relative density of pairs of individuals separated by distance *r*, normalized against CSR. By definition, *g*(*r*) = 1 under CSR, values *g*(*r*) *>* 1 indicate aggregation, and values *g*(*r*) *<* 1 indicate overdispersion (regular spacing). In analyzing the simulations, I treat each individual as located at the center of its occupied grid cell, so that the community can be regarded as a point pattern in continuous space with unit cell spacing. I consider *r* in units of grid cell lengths. Analysis of *g*(*r*) allows quantification of aggregation at multiple spatial scales. This was implemented using the R package spatstat (Baddeley *et al*., 2015).

### Analysis and simulations

I combined mathematical analyses with simulations of the spatially explicit model. First, I used invasion analysis (Turelli, 1978), deriving an approximate analytical condition under which a rare species deterministically increases in abundance when introduced into a community of *N* resident species in the local community. This was accomplished by deriving when the quantity 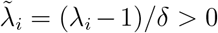 as *p*_*i*_ → 0 where 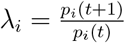 is the finite rate of increase and *δ* is the adult mortality rate. I refer to 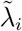 as the weighted growth rate of species *i* (Chesson, 2003). See Supplementary Information “Invasion criteria” for detailed derivations.

Next, I conducted forward-in-time simulations on the 175×175 grid with periodic boundary conditions (a torus) to avoid hard-edge boundary artifacts (missing neighbors and truncated dispersal) in local interactions and dispersal. Within each parameter sweep, the habitat realization (and random seed) was held fixed for comparability; due to computational constraints, each parameter combination is represented by a single simulation run. The local community was initialized with species drawn from the regional pool. In most simulations, I assumed the regional pool contained *L* = 500 species, although I also tested how the regional pool size affects local community species richness. Simulations were run for ∼ 10,000 generations (where one generation corresponds to ∼ 1*/δ* time steps) — sufficient for species richness to reach a dynamic equilibrium maintained by the balance of extinction and immigration (see Supplementary Information “Confirmation of dynamic equilibrium”).

With these specifications, I conducted four main sets of simulations. **(1) Regional pool size and local species richness**. I performed simulations varying the regional pool size under five different scenarios: when only JC-effects occur, when only habitat partitioning occurs, and when both occur under low, intermediate, and high levels of spatial autocorrelation (defined by the magnitude of the nugget). **(2) Mechanism synergy analysis**. Using the parameterizations described in above section, I examined the *synergy* of JC-effect – habitat partitioning interactions. I defined the synergy of JC-effects and habitat partitioning as the factor by which species richness under their joint operation exceeds the sum of their separate contributions. For example, with *M* = 7 and *σ*_*h*_ = 0.02, letting “SR” denote species richness:

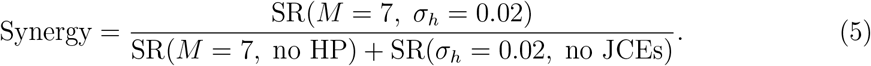

I refer to JC-effect–habitat partitioning interactions with synergy *>* 1 as “synergistic” and those with *<* 1 as “antagonistic”. I ran simulations to examine how the level of spatial auto-correlation and dispersal limitation modifies the JC-HP synergy. **(3) Drivers of aggregation and coexistence**. I investigated how processes shaping species’ spatial distributions — dispersal limitation (*σ*_*h*_), JC-effect strength (*a*), and spatial autocorrelation (quantified by the nugget) — combine to influence the relationship between aggregation and species richness. **(4) The spatial scale of JC-effects**. I examined how the spatial scale of JC-effects *per se* influences species richness. The overall strength of JC-effects depends on both their baseline intensity (*a*) and their spatial scale (*M*), such that the total JC-pressure generated by an individual tree is *aM*^2^ (as defined above). To isolate the role of JC-effect spatial scale, I varied *M* while holding *aM*^2^ = *K* constant, including the “global” JC-effect limit of 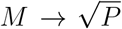 (where *P* is the total number of patches), in which every patch experiences JC-effects from every adult. I then compared how species richness responds to JC-effect spatial scale when (1) only JC-effects operate (with and without dispersal limitation) and (2) when both JC-effects and habitat partitioning are present.

All simulations were implemented in Julia version 1.11 (Bezanson *et al*., 2017). Figures were produced in R version 4.3.2 (R Core Team, 2021).

## Results

### Invasion criteria

This invasion analysis has two aims: (i) to quantify how the contributions of JC-effects, habitat partitioning, and their interaction scale with *N*, the number of resident species in the local community, and (ii) to distinguish how aggregation arising from habitat specialization versus dispersal limitation alters the contribution of JC-effects to coexistence. Accordingly, I consider two cases: (1) JC-effects with habitat partitioning under global dispersal, and (2) JC-effects alone under dispersal limitation.

#### JC-effects, habitat partitioning, and global dispersal

Suppose species *i* is a rare invader with intrinsic fitness *Y*_*i*_, habitat optimum *h*_opt,*i*_, and assume dispersal is global. Let 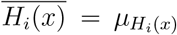 denote the mean performance of species *i* across space *x*; here and below, spatial averages (e.g., 𝔼_*x*_[·]) are taken at the resident community’s long-term equilibrium. Then species *i* invades if its weighted growth rate is positive:

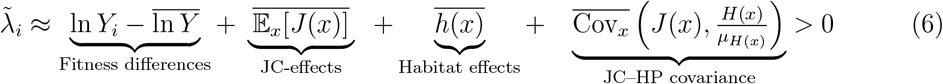

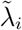 consists of four terms: a fitness difference term and three terms capture how JC-effects, habitat effects/partitioning, and their interaction contribute to invasion. I discuss each term in detail below. Note that this invasion criterion is flexible to different functional forms of *J*(*x*) and *H*(*x*). Below I emphasize the functional forms presented in the Methods section, and assume species have identical breadths, 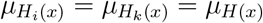 (*k* = 1, 2, …, *N, k* ≠ *i*), for analytical tractability. The below approximations are accurate within the range of parameters utilized in simulations shown in the main text (Supplementary Information “Analysis of approximations” for details).

First, consider fitness differences. 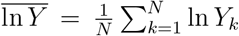 is the mean log fitness of the resident species. Thus, ln 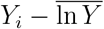 represents the invader’s fitness difference relative to the mean resident fitness. If this difference is negative, the contributions of JC-effects, habitat effects, and the JC-HP covariance must be sufficiently strong to permit invasion.

Second, JC-effects contribute to invasion. The direct contribution of JC-effects to co-existence, 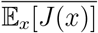, is the mean JC-effect strength across space *x*, averaged over resident species: 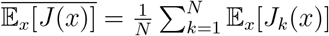. Using the form in equation 2, this yields

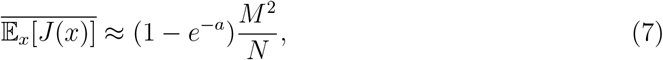

where *a* is the baseline JC-effect strength, *M* is the spatial scale, and *N* is the number of resident species. This approximation assumes adults are randomly distributed in space. Intuitively, stronger baseline JC-effects (*a*) and larger neighborhoods (*M*) promote coexistence. However, because 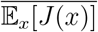 decreases in proportion to 1*/N*, the ability of JC-effects to maintain coexistence declines rapidly as species richness increases (Fig. 2A, blue bars).

**Figure 2:**
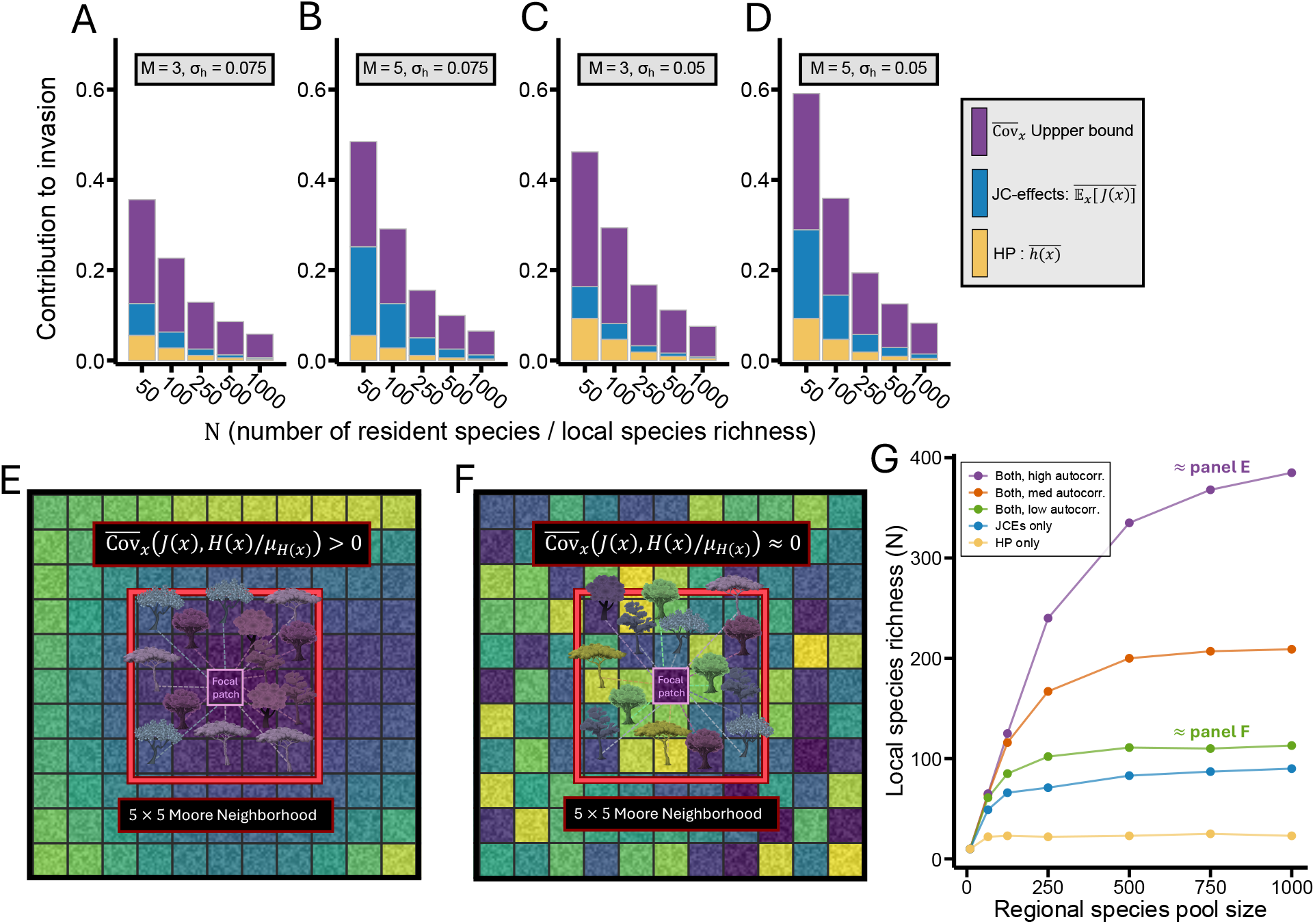
The JC–HP covariance can strongly promote invasion in highly diverse communities. (A-D) Analytical results showing the contributions to coexistence from habitat partitioning (HP; yellow), JC-effects (blue), and the upper bound of the JC–HP covariance (purple) as a function of resident species richness *N*. All plots use *a* = 0.5. (E) and (F) Illustrate JC–HP covariance when habitats are strongly vs. weakly spatially autocorrelated, respectively. Color indexes tree habitat preference. (G) Simulations. The x-axis shows the number of species in the regional pool, and the y-axis shows local species richness *N* at equilibrium. Colored curves correspond to five scenarios labeled in the legend. In all simulations, *σ*_*D*_ = ∞, *a* = 0.5, *ν* = .3 × 10^−5^, *Y* ∼ lognormal(*µ* = 0, *σ*_*Y*_ = 0.3), *M* = 7 (except HP only), *σ*_*h*_ = 0.02 (except JCEs only), and habitat range = 0.5. Low, medium, and high autocorrelation are defined by nugget = 10 (similar to panel F), 0.1, or 0 (similar to panel E), respectively.

Aggregation reduces 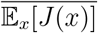. To see this, I derived an alternative approximation that relaxes the assumption of spatial randomness:

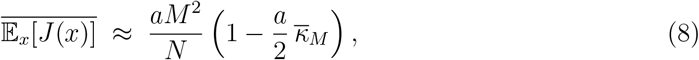

where 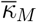 is the mean intraspecific aggregation of residents at the *M* × *M* scale (equation 4). This approximation is valid when *a* and 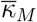 are not very large (i.e., the expression remains non-negative within valid parameter space). Thus, aggregation *per se* (larger 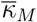) reduces the ability of JC-effects to promote invasion (see Supplementary Information, “Moment-based approximation of 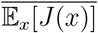” for derivation). This result is a direct consequence of the non-linear nature of *J*(*x*), as described by Jensen’s inequality (Jensen, 1906).

Third, habitat effects contribute to invasion. The “Habitat effects” term 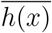 captures the contribution to invasion that arises from differences in species’ responses to the environment. Conceptually, this term quantifies the extent to which a rare invader’s habitat response *H*_*i*_(*x*) overlaps the resident species’ habitat responses relative to how much residents overlap with themselves and each other. A full derivation is given in the Supplementary Information (“Derivation of 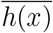”), but the structure is particularly transparent when all species (including the invader) have the same average correlation in their habitat responses. In this case

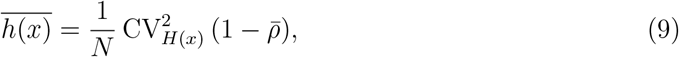

where 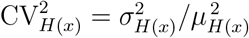 is the squared coefficient of variation of habitat responses, and 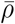 is the average correlation in habitat responses across species. Note: *σ*_*H*(*x*)_ ≠ *σ*_*h*_. Intuitively, habitat effects are strongest when responses are weakly correlated 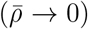, and it vanishes as responses become more tightly aligned 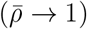.

Using the Gaussian response function from equation (1) and assuming resident optima are randomly distributed (i.e., *h*_opt_ ∼ Unif[0, 1]), one can obtain a simple closed-form approximation in the regime of specialized niches (*µ*_*H*(*x*)_ ≪ 1). In this limit, 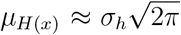 and 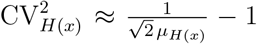. Substituting into equation (9) and taking 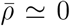 (as expected when habitat optima are randomly distributed) yields

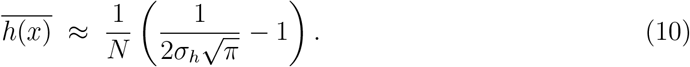

Thus, narrower niches (smaller *σ*_*h*_) strengthen the contribution of habitat partitioning to invasion, but — just as with JC-effects — the contribution to invasion is inversely proportional to *N* (Fig. 2A, yellow bars). Conceptually, as *N* increases, the resident community more completely samples the habitat axis, reducing the invader’s expected habitat-mediated advantage. If resident habitat optima are evenly spaced rather than randomly distributed, 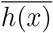 can decay faster than ∝ 1*/N* with increasing *N* (Supplementary Information “Derivation of 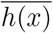)”).

Fourth, the interaction between JC-effects and habitat partitioning (the JC–HP covariance) is given by

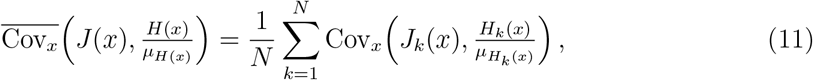

where Cov_*x*_(*J*_*k*_(*x*), *H*_*k*_(*x*)) is the spatial covariance between JC-effect strength and habitat effects of species *k*. A positive JC–HP covariance promotes invasion. The mechanism reflects a key principle: invaders experience their strongest competition in habitats favorable to them, which are also preferred by resident species with similar optima. If those residents disproportionately generate JC-effects in those same habitats – as occurs under spatial autocorrelation – resident fitness is reduced more strongly than if JC-effects were induced in unfavorable habitats. This yields 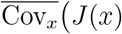, *H*(*x*)*/µ*_*H*(*x*)_ *>* 0 (Fig. 2E). In contrast, when habitats are randomly distributed, 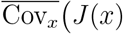, *H*(*x*)*/µ*_*H*(*x*)_ ≈ 0, and JC-effects and HP do not (or weakly) interact (Fig. 2F).

The JC–HP covariance term can exceed the direct effects of 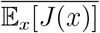 and 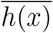. Using the Cauchy–Schwarz inequality (Cov (*J*(*x*), *H*(*x*)) ≤ *σ*_*J*(*x*)_*σ*_*H*(*x*)_), I derived two approximate upper bounds. Conceptually, the bound occurs when species’ JC-effects are maximally concentrated in habitats favorable to them (Fig. 2E; see Supplementary Information, “Upper bound on JC–HP covariance”).

First, heuristically,

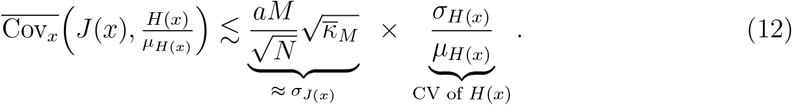

The JC–HP covariance – which promotes invasion – increases with mean intraspecific aggregation 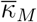. This contrasts with the direct JC-effect term, which decreases in strength with aggregation (eq. 8).

Second, assuming *µ*_*H*(*x*)_ ≪ 1 and species are maximally aggregated in space (so that *σ*_*J*(*x*)_ attains its upper bound), a tighter approximation is

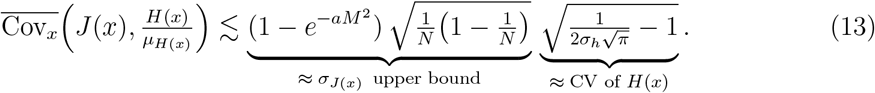

For both bounds, the contribution of the JC–HP covariance to invasion is 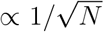, which decays more slowly with species richness than the direct effects of JC-effects or habitat partitioning (∝ 1*/N*). Thus, even when 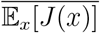 and 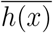 are small and contribute little to invasion — which occurs when *N* is large — the JC-HP covariance can be large (Fig. 2A–D; purple vs. blue and yellow bars). Note that Fig. 2 shows equation (13).

#### JC-effects and dispersal limitation

With local dispersal, propagules deposited near a parent tree are subject to its JC-effects (Fig. 1D, i). Let *d* denote the proportion of propagules that disperse far enough to avoid JC-effects within the parent’s *M* ×*M* Moore neighborhood. In this case, to a first approximation, invasion occurs when

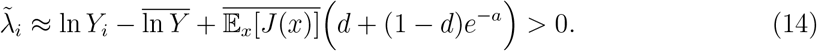

See Supplementary Information (“JC-effects, dispersal limitation, and no habitat partitioning”) for the derivation. Dispersal limitation reduces the contribution of JC-effects to invasion relative to global dispersal through two mechanisms.

First, the loss of locally dispersed propagules weakens the impact JC-effects have on invasion. The term 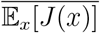 is multiplied by the factor *d* + (1 − *d*)*e*^−*a*^ ≤ 1 and therefore declines as dispersal becomes more limited (smaller *d*). This is because a fraction 1 − *d* of propagules remain near the parent and experience JC-effects, weakening the advantage of rarity.

Second, dispersal limitation leads to intraspecific aggregation (Fig. 1D, iii). As shown in equation 8, aggregation directly reduces 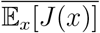.

### Simulation results

#### JC-effects and habitat partitioning in isolation

Local species richness *N* maintained by JC-effects alone or habitat partitioning alone saturated rapidly with regional pool size (Fig. 2G, blue and yellow curves).

When only JC-effects operated, the number of coexisting species increased with their spatial scale (*M*). With *a* = 0.5, up to ∼100 species persisted (Fig. 3A, yellow points, *M* = 9; Fig. 2G, blue points). Dispersal limitation (*σ*_*D*_ = 1.5) reduced richness relative to global dispersal, sometimes by more than 50% (compare the yellow points in in Fig. 3A and Fig. 3C; also see Fig. 5B).

**Figure 3:**
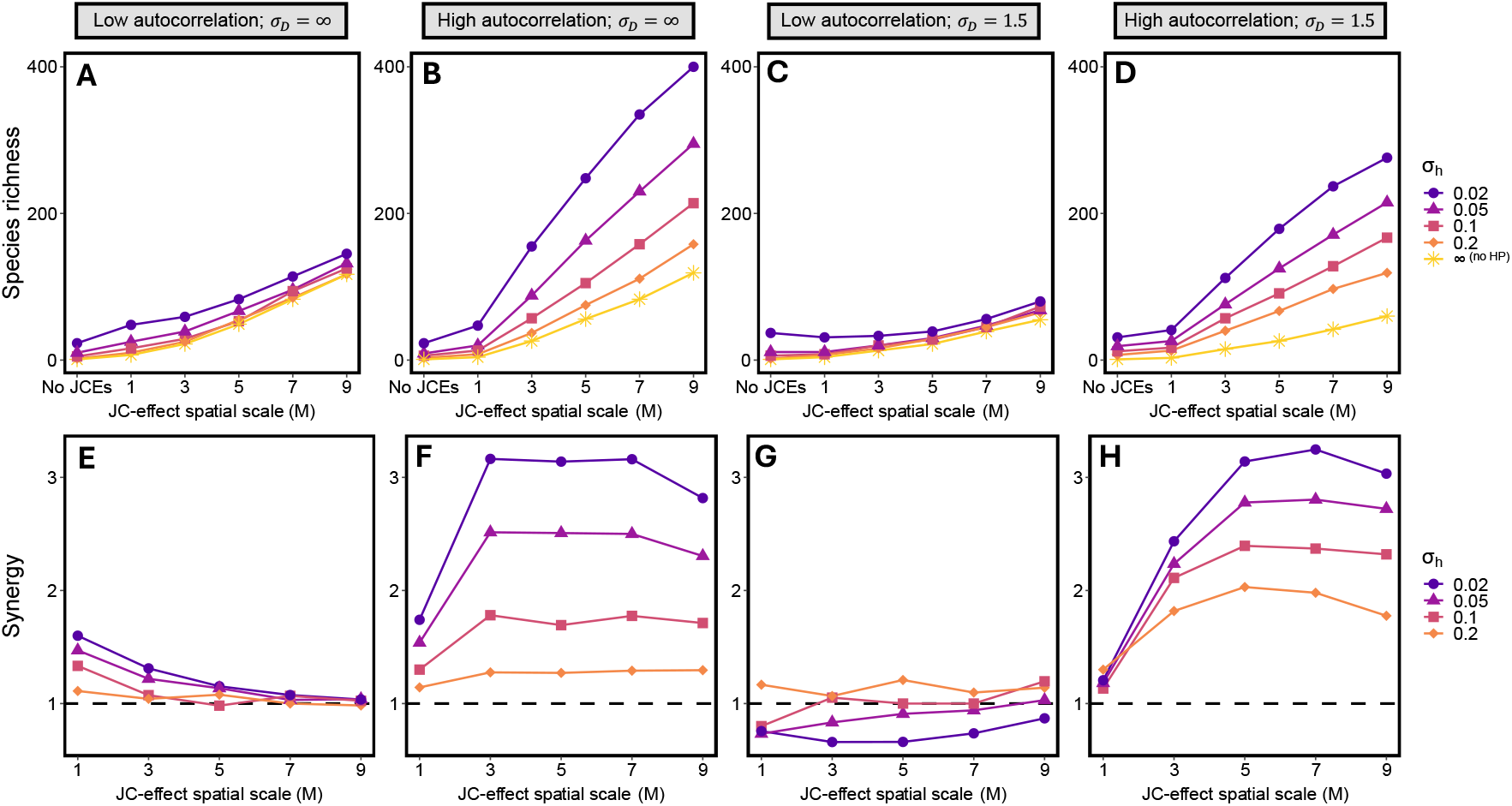
Spatial autocorrelation and dispersal limitation jointly influence JC–HP synergy. Each column represents a different case defined by the level of spatial autocorrelation and dispersal limitation (see panel labels). (A–D) Simulation outputs, showing equilibrium species richness (y-axis) across combinations of JC-effect scale (*M*, x-axis) and niche breadth (*σ*_*h*_, points/color scale). (E–H) show the corresponding synergy for each parameter combination from the simulations depicted in the above plots. In all simulations, *a* = 0.5, *ν* = 0.3 × 10^−5^, *Y* ∼ lognormal(*µ* = 0, *σ*_*Y*_ = 0.3), *δ* = 0.2, regional pool size: *L* = 500, and the habitat range was set to 0.5. High spatial autocorrelation corresponds to nugget = 0, while low autocorrelation corresponds to nugget = 10.

When only habitat partitioning operated, richness increased with greater specialization (smaller *σ*_*h*_; Fig. 3, “No JCEs”). Roughly 20–30 species persisted under narrow niches and global dispersal (*σ*_*h*_ = 0.02; see Fig. 1A, purple curve, for niche width). Unlike JC-effects, habitat partitioning yielded higher richness under dispersal limitation (“No JCEs” in Fig. 3A,B vs. C,D), with ∼30–40 species coexisting under the narrowest niches examined (Fig. 3C,D).

These results should be interpreted as a frame of reference: larger *a* or *M*^2^ and smaller *σ*_*h*_ would all increase species richness. Below, I examine how the joint operation of JC-effects and habitat partitioning differs from the sum of their parts relative to these baseline results.

#### Joint operation of JC-effects and habitat partitioning

When both mechanisms acted together, outcomes depended strongly on the degree of spatial autocorrelation in habitat structure. Local species richness *N* saturated quickly to a maximum level as a function of the size of the regional pool under low spatial autocorrelation (Fig. 2G, green curve). In contrast, saturation was much slower under moderate or strong autocorrelation (blue and purple curves, respectively). For example, under otherwise identical parameters, highly autocorrelated environments maintained ∼400 species, compared with ∼100 species in weakly autocorrelated environments (Fig. 2G, purple vs. green curves at a regional pool size of 1000). More generally, species richness was higher under high spatial autocorrelation (compare Fig. 3A to 3B and 3C to 3D) and under global dispersal (compare Fig. 3A to 3C and 3B to 3D) when both JC-effects and habitat partitioning operated.

JC–HP interactions ranged from weakly synergistic to antagonistic when spatial autocorrelation was low, but were typically highly synergistic when autocorrelation was high (Fig. 3; see equation 5 for the definition of synergy). Under low autocorrelation and global dispersal (*σ*_*D*_ = ∞; Fig. 3A,E), JC-effects and habitat partitioning were moderately synergistic when *M* = 1, with synergy up to ≈ 1.5 under narrow niches (*σ*_*h*_ = 0.02). However, synergy declined with JC-effect scale *M*, disappearing by *M* = 7 (Fig. 3E). With dispersal limitation (*σ*_*D*_ = 1.5; Fig. 3C,G), synergy was largely absent; JC-HP interactions were antagonistic under narrow niches (small *σ*_*h*_; Fig. 3G, orange and green curves).

High spatial autocorrelation yielded synergistic interactions in all cases. While large *M* eliminated synergy under low spatial autocorrelation (Fig. 3E), it amplified synergy under high autocorrelation, with values exceeding synergy *>* 3 (Fig. 3F,H). Moreover, synergy was often greater under dispersal limitation than under global dispersal (compare Fig. 3F to 3H), again in contrast to the low-autocorrelation case, where dispersal limitation led to antagonistic interactions.

The increase in species richness associated with positive spatial autocorrelation is driven by the JC–HP covariance. In simulations varying the range and nugget of spatial variation, species richness closely tracked the JC–HP covariance (compare Fig. 4A,B). Richness increased approximately linearly with the JC–HP covariance term (Fig. 4C). This linear association was robust to when species differed in baseline JC-effect strength *a* (Supplementary Information, “Interspecific variation in baseline JC-effect strength”) and when species exhibited a specialist–generalist trade-off in their habitat responses (Supplementary Information, “Tolerance–fecundity trade-off model”).

**Figure 4:**
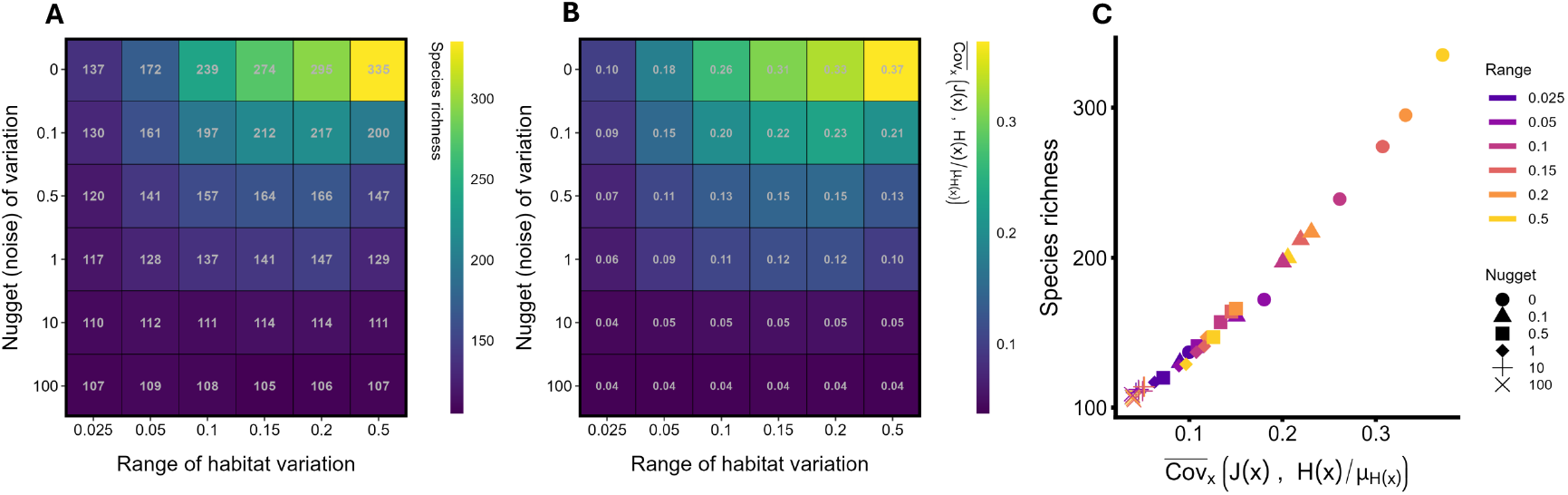
When both JC-effects and habitat partitioning operate, species richness increases with abiotic habitat spatial autocorrelation, driven by the JC–HP co-variance. All panels depict simulations in which the range and nugget (noise term) of spatial variation were varied. (A) and (B) show how the range and nugget influence species richness and the JC–HP covariance, respectively. (C) shows the relationship between species richness and the JC–HP covariance across simulations. Color indexes the range, shape indexes the nugget. The JC-HP covariance was numerically calculated from the spatial distribution of habitats and adults in the landscape at the end of simulations. Parameters: *a* = 0.5, *ν* = 0.3 × 10^−5^, *δ* = 0.2, *σ*_*h*_ = 0.02, *Y* ∼ lognormal(*µ* = 0, *σ*_*Y*_ = 0.3), *M* = 7, and dispersal is global.

#### Intraspecific aggregation and species richness

The relationship between species aggregation and species richness differed qualitatively depending on the mechanism generating aggregation. Consider first the case of JC-effects alone. The direct impact of JC-effects on invasion increases with the baseline strength *a* (equations 7 and 8), and correspondingly, species richness increases with *a* (Fig. 5A). When *a* is large (e.g., *a >* 10), conspecific recruitment is effectively impossible within the Moore neighborhood. This reduces aggregation and produces regular/overdispersed conspecific spacing (Fig. 5M; 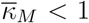 in Fig. 5E; pair-correlation function *g*(*r*) *<* 1 for small *r* in Fig. 5M). Thus, species richness decreased with increasing aggregation (Fig. 5F). Similarly, dispersal limitation reduced richness with JC-effects operated alone (Fig. 5), again generating a negative relationship between richness and aggregation (Fig. 5F). In simulations with modest JC-effect strength (*a* = 0.5), dispersal limitation often overcame the repulsion of JC-effects (e.g., *g*(*r*) *>* 1 for small *r* in Fig. 5J,N), which resulted in low species richness. In sum, under JC-effects alone, aggregation and richness were negatively associated.

**Figure 5:**
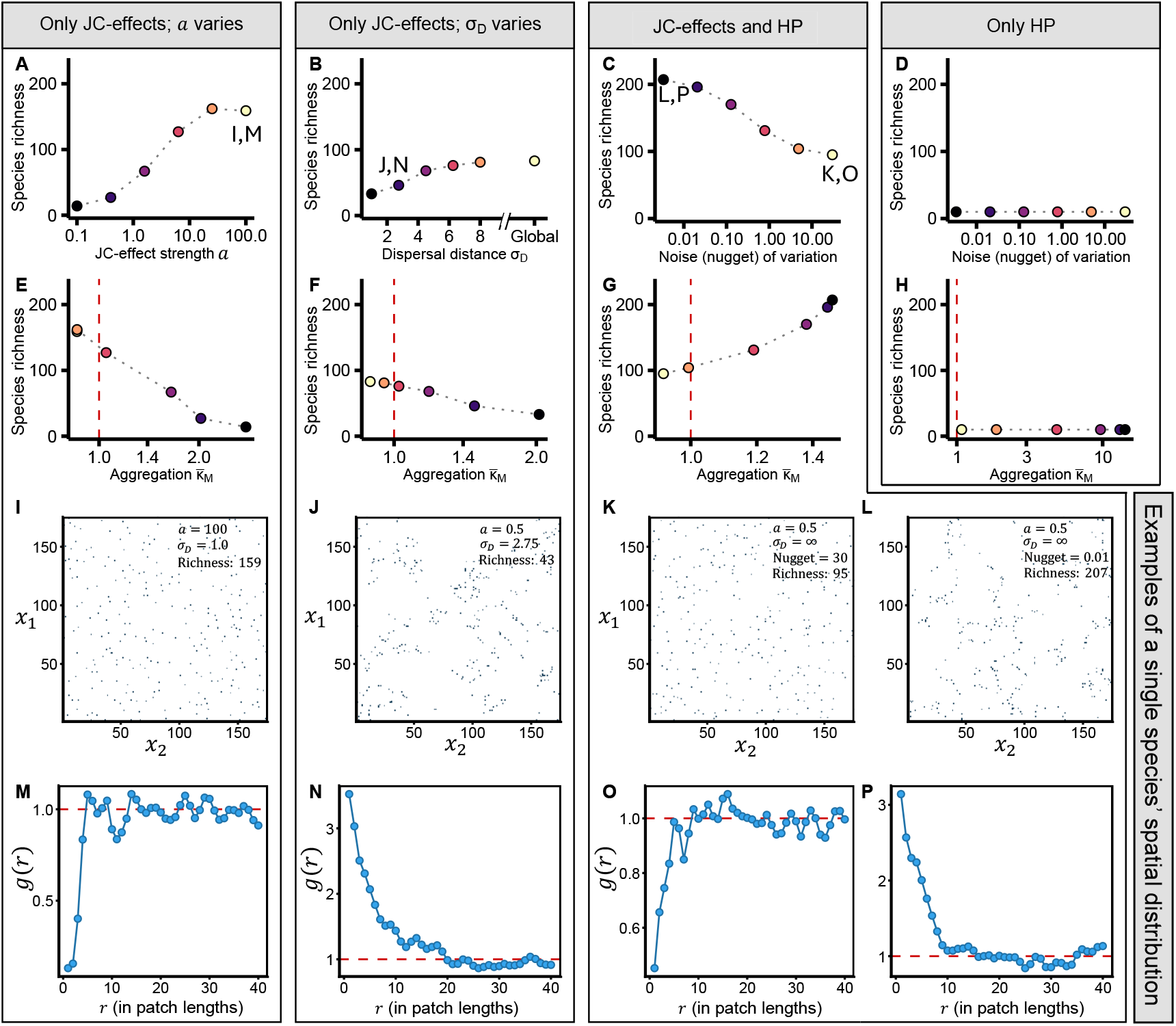
Aggregation driven by habitat partitioning vs. dispersal limitation has qualitatively different effects on species richness. (A–D) show species richness from simulations as a function of the specified parameters; (E–H) show richness as a function of mean intraspecific aggregation 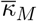 in the corresponding simulations in the above panel (within the same rectangular box). Points of the same color in, e.g., (A) and (E) are from the same simulation. (A,E) JC-effects only, varying *a* (baseline JC-effect strength); *σ*_*D*_ = 1.0. (B,F) JC-effects only, varying dispersal limitation *σ*_*D*_; *a* = 0.5. (C,G) Joint operation of JC-effects and habitat partitioning, varying the nugget of habitat variation; *σ*_*D*_ = ∞. (D,H) Same parameters as (C,G), but with habitat partitioning only. (I–L) Spatial distribution of one species with an abundance of ∼300 individuals, selected from a labeled simulation in A, B, or C. Note the plots show two examples from the “JC-effects and HP” case (K, L) and no examples from the “Only HP”. (M–P) Pair-correlation function *g*(*r*) for the focal individual in the above panel. *g*(*r*) *>* 1 and *g*(*r*) *<* 1 indicate aggregation and repulsion, respectively. *r* indicates the number of patch lengths. All simulations use *ν* = 0.3 × 10^−5^, *Y* ∼ lognormal(*µ* = 0, *σ*_*Y*_ = 0.3), *δ* = 0.2, *M* = 7, *σ*_*h*_ = 0.05, and habitat range = 0.5 (when relevant).

In contrast, when both JC-effects and habitat partitioning were present, aggregation driven by habitat specialization was associated with greater species richness. As in Fig. 4, richness increased as the nugget (environmental noise) decreased (Fig. 5C). This produced a positive relationship between richness and aggregation (Fig. 5G): low richness occurred when species exhibited regular spacing (*g*(*r*) *<* 1 for small *r*; Fig. 5O), whereas high richness occurred when species were highly aggregated at small spatial scales (*g*(*r*) *>* 1 for small *r*; Fig. 5P).

The relationship between richness and aggregation in Fig. 5G does not arise from habitat partitioning *per se*. Under global dispersal, aggregation driven by habitat specialization was unrelated to species richness when JC-effects were absent (Fig. 5D,H). Thus, the positive aggregation–richness association stems entirely from how habitat partitioning modifies the operation of JC-effects.

#### The spatial scale of JC-effects and species richness

The relationship between JC-effect spatial scale (*M*) and species richness depended on whether JC-effects acted alone or alongside habitat partitioning, in simulations with fixed total JC-effect pressure (*aM*^2^ = *K*). With JC-effects alone, richness increased with *M* and plateaued under global JC-effects (Fig. 6A). Across finite *M*, richness was higher under effectively global propagule dispersal (*σ*_*D*_ = ∞) than under propagule dispersal limitation (*σ*_*D*_ = 1), but the two treatments converged under global JC-effects (Fig. 6A). These patterns follow directly from Eq. 8. Holding *aM*^2^ = *K* constant, the invasion contribution from JC-effects to invasion scales approximately as

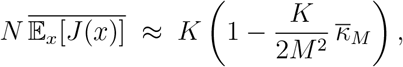

which is a strictly increasing function of *M* (Fig. 6B). Propagule dispersal limitation increases 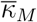, reducing 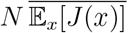 (Fig. 6B, gray circles vs. red triangles). As *M* → ∞, the 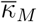 term vanishes, causing 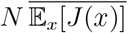 to converge for *σ*_*D*_ = ∞ and *σ*_*D*_ = 1 (Fig. 6B, “Global”).

**Figure 6:**
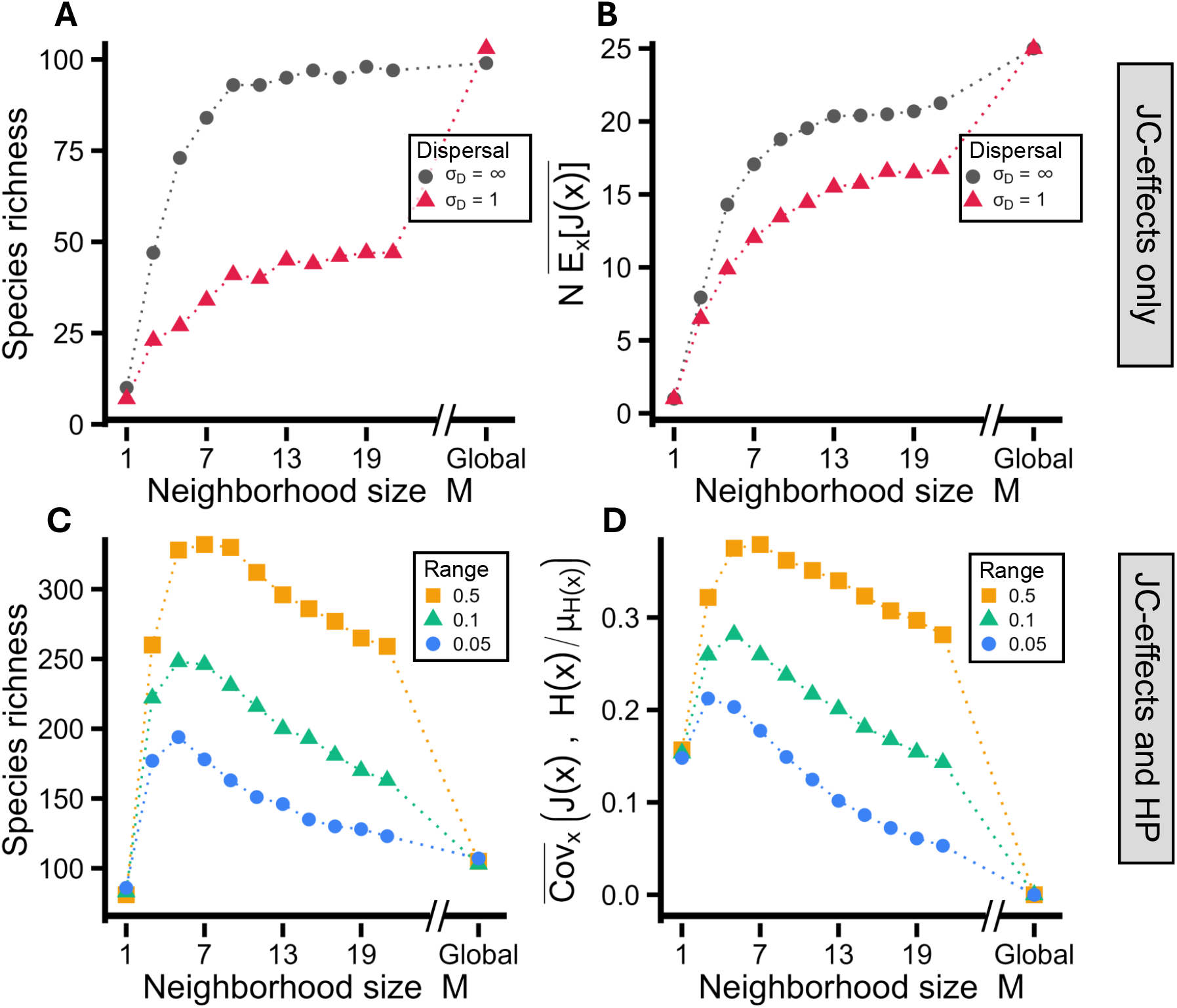
The relationship between the spatial scale of JC-effects and species richness differs depending on whether JC–HP covariance is present. (A) Species richness as a function of *M* (neighborhood size / spatial scale) with JC-effects alone under global dispersal (*σ*_*D*_ = ∞) and strong dispersal limitation (*σ*_*D*_ = 1). (B) depicts analytical results, showing the quantity 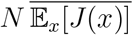 as a function of *M* ; mean JC-effect strength 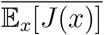 was numerically calculated from the spatial distribution of adults in the landscape at the end of simulations. (C) Species richness as a function of *M* when JC-effects and habitat partitioning operate, for three habitat range values (nugget = 0). (D) JC–HP co-variance as a function of *M*. The JC-HP covariance was numerically calculated from the spatial distribution of habitats and adults in the landscape at the end of simulations. In simulations, JC-effect pressure was held constant (*aM*^2^ = *K* = 25). All simulations used *ν* = 0.3 × 10^−5^, *Y* ∼ lognormal(*µ* = 0, *σ*_*Y*_ = 0.3), *δ* = 0.2; in (C,D), *σ*_*h*_ = 0.02.

By contrast, species richness peaked at an intermediate spatial scale *M* when both JC-effects and habitat partitioning operated (assuming habitats are spatially autocorrelated; Fig. 6C). This occurs because the JC–HP covariance also peaked at intermediate *M* (Fig. 6D). When JC-effects extend over large spatial scales (*M >* 5 or *M >* 7; Fig. 6C), they spill over into relatively poor habitat types, reducing the JC–HP covariance. This decline is slower when spatial autocorrelation has a relatively large range of variation (Fig. 6C–D; compare orange to green and blue points).

## Discussion

Quantifying how coexistence mechanisms interact is crucial for understanding species richness. I show that two leading explanations for tropical forest diversity — Janzen–Connell (JC) effects and habitat partitioning (HP) — can interact in ways that fundamentally alter their implications for coexistence. When habitats are positively spatially autocorrelated, habitat specialization clusters species in favorable environments and JC and HP act synergistically, promoting high richness. By contrast, aggregation generated by dispersal limitation weakens the ability of JC-effects to maintain coexistence. I capture these outcomes with a new metric — the JC–HP covariance — which quantifies the extent to which JC-effects are concentrated in favorable versus unfavorable environments.

### Conceptual and theoretical context

JC-effects and habitat partitioning each face limitations when operating alone. JC-effects generate negative frequency dependence by reducing recruitment near conspecific adults, but their contribution to invasion declines with species richness (∝ 1*/N*) (also see Chisholm & Fung, 2020; Smith, 2022). Habitat partitioning allows species with different habitat preferences to coexist (Snyder & Chesson, 2003; Wright, 2002), yet its effect is also inversely proportional to *N* and cannot sustain high richness unless niches are quite narrow.

When the two mechanisms act together, their interactions can greatly exceed the sum of their parts. In spatially autocorrelated habitats, species cluster in environments where they perform best, and resident adults disproportionately impose JC-effects on those same favorable locations. Invaders that share habitat preferences therefore encounter weakened competitors, because residents are heavily suppressed precisely where they would otherwise compete the strongest. The strength of this interaction is captured by the JC–HP covariance, which may decay more slowly with species richness 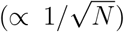 than the direct effect of either mechanism (∝ 1*/N*), and so remains strong even in highly diverse communities. This allows habitat partitioning that might otherwise sustain only 10–30 species to maintain hundreds when coupled with JC-effects (Figs. 2G, 3B). While extreme parameter choices could in principle allow high richness under JC-effects alone (very large *a* and *M*) or habitat partitioning alone (very small *σ*_*h*_), JC–HP interactions achieve the same outcome under far less restrictive conditions because their contribution to invasion erodes much more slowly with *N*.

This result both builds on and contrasts with previous theory. Stump & Chesson (2015) analyzed a spatially implicit model in which JC-effects operate only at the parent’s patch, and found little interaction between JC-effects and habitat partitioning. However, when spatial structure is modeled explicitly, I find that JC-effects and habitat partitioning interact strongly via the JC–HP covariance. Stump & Chesson (2015) also argued that distance-dependent specialized predation is less able to stabilize coexistence than globally acting enemies; I recover this when only JC-effects operate. With habitat partitioning, however, the conclusion reverses in spatially explicit settings: local, distance-dependent JC-effects maintain higher richness than global JC-effects. This reversal occurs because global JC-effects (i.e., globally dispersing natural enemies) do not generate a JC–HP covariance, whereas local effects do. Overall, understanding JC-effects and their interactions with other coexistence mechanisms requires spatially explicit models.

JC–HP interactions can also be antagonistic. When habitat partitioning acts alone, dispersal limitation promotes coexistence because rare species disproportionately recruit into favorable habitats, generating a fitness–density covariance (Chesson, 2000a; Snyder, 2008). JC-effects can erode this fitness–density covariance by reducing conspecific recruitment in those favorable patches, likely leading to the observed antagonistic interactions. Notably, interactions were antagonistic only under low autocorrelation (when habitat structure did not generate a JC–HP covariance) and when trees were dispersal limited. Thus, whether JC–HP interactions are synergistic or antagonistic depends qualitatively on both habitat structure and dispersal.

### Aggregation, JC-effects, and coexistence

The results of this study suggest the need to reinterpret the relationship between intraspecific aggregation, JC-effects, and coexistence. Overdispersion is often taken as evidence of strong JC-effects (Janzen, 1970; Chave *et al*., 2002; Wiegand & Moloney, 2013), whereas the pervasive aggregation of tropical trees has been used to question their efficacy. For example, Chisholm & Fung (2020) argue that the close proximity of conspecific adults in tropical forests makes it implausible that JC-effects are strong enough to prevent local recruitment, and therefore unlikely to maintain high species richness. Consistent with this view, I find that JC-effects alone maintain substantial richness only when baseline strength *a* is very large; such values eliminate recruitment near adults and generate pronounced overdispersion, in contrast with the near-universal aggregation of tropical trees (Condit *et al*., 2000). However, when habitat partitioning operates alongside JC-effects, species richness can increase with intraspecific aggregation via the JC–HP covariance. Accordingly, findings that habitat partitioning and dispersal limitation explain spatial distributions more than JC-effects (e.g., Zhu *et al*., 2013; Barry & Schnitzer, 2021) are not inconsistent with JC-effects contributing substantially to diversity maintenance (also see Kalyuzhny *et al*., 2023). Taken together, JC–HP interactions provide a more empirically plausible pathway to high diversity maintenance than JC-effects alone once spatial aggregation is taken into account.

Aggregation is not inherently indicative of strong or weak diversity maintenance by JC-effects; its consequences depend on the processes that generate it. Habitat specialization promotes coexistence by concentrating JC-effects in favorable environments, whereas dispersal limitation undermines coexistence by reducing the average strength of JC-effects over space and preventing rare species’ propagules from escaping conspecific effects. This result is consistent with relatively recent theory (Stump & Chesson, 2015; Chisholm & Fung, 2020) although it contrasts with earlier work (Adler & Muller-Landau, 2005). If dispersal limitation overrides the repulsion generated by JC-effects and habitat partitioning does not operate, it likely signals that JC-effects are weak and cannot maintain high richness. In contrast, JC-effects too weak to prevent aggregation from habitat specialization can sustain high richness through the JC–HP covariance. However, notably, dispersal-based aggregation that decouples recruits from parent locations can enhance coexistence (e.g. via animal dispersal; Wiegand *et al*., 2021, 2025); this form of dispersal limitation may generate different JC-effect – dispersal limitation interactions.

Documenting aggregation alone is thus insufficient for inferring the importance of JC-effects. Dispersal limitation and habitat specialization can generate superficially similar aggregation patterns (Fig. 5J,N vs. L,P) but have qualitatively different implications for how JC-effects regulate species richness. Identifying the processes that generate aggregation is therefore essential to assess how spatial patterns relate to the efficacy of density-dependent processes such as JC-effects.

### Empirical considerations for forest communities

These theoretical results highlight a promising empirical target: directly measuring the JC–HP covariance in forest plots. The importance of habitat heterogeneity varies among sites (Shen *et al*., 2013), and the strength of its interaction with local density dependence likely varies as well. For example, the BCI forest dynamics plot in Panama was deliberately chosen for relatively low spatial heterogeneity (Hubbell *et al*., 2024), whereas the Lambir Hills plot in Malaysia exhibits sharp edaphic contrasts, steep species turnover, and harbors roughly four times as many tree species in a comparable area (52 ha vs. 50 ha; Palmiotto *et al*., 2004; Russo *et al*., 2005). Stabilizing conspecific negative density dependence has been detected at both sites (Hülsmann *et al*., 2024). Lambir Hills’ higher richness may therefore reflect not habitat heterogeneity *per se*, but the way heterogeneity interacts with density-dependent processes such as JC-effects.

A key challenge is to quantify how habitat heterogeneity shapes the relative importance of JC-effects, habitat partitioning, and their interaction. Doing so requires not only estimating the strength of each mechanism, but also distinguishing whether conspecific aggregation arises primarily from dispersal or from habitat association. Targeted approaches that separate dispersal-dependent from habitat-linked clustering are therefore essential. One direction is to combine generalized linear mixed models (GLMMs) with spatial point-pattern analyses to estimate the JC–HP covariance from censused forest plot data. Using repeatedly censused seedling or sapling performance data at mapped locations (e.g., Comita *et al*., 2023), GLMMs can quantify how conspecific (and heterospecific) neighborhood covariates and abiotic gradients (e.g., topography, elevation, soils) jointly influence survival or growth (e.g., Yao *et al*., 2020; Huang *et al*., 2022; Wiegand & Moloney, 2013; Wiegand *et al*., 2021). From the fitted model, one can construct (i) a habitat suitability surface *H*(*x*) (predicted baseline performance across space, holding neighborhood density fixed) and (ii) a JC-effect surface *J*(*x*), defined as the fitted change in survival at location *x* attributable to conspecific neighbors (i.e., the predicted survival difference with the observed conspecific neighborhood covariate(s) at *x* versus setting those covariate(s) to zero, holding other predictors fixed). The JC–HP covariance can then be directly estimated as Cov(*J*(*x*), *H*(*x*)). In practice, *H*(*x*) can be defined using the best-supported habitat representation for a given system (continuous edaphic/topographic covariates, ordinated environmental axes, or discrete habitat classes, e.g. the habitat categories commonly used at BCI; Harms, 2024). Computing the covariance under alternative habitat definitions provides a straightforward way to assess which abiotic variables most strongly shape JC–HP coupling.

Experimental manipulations provide a complementary approach to empirically measure the JC–HP covariance. Factorial designs that cross conspecific proximity with fungicide treatments have already demonstrated the role of specialized pathogens in generating con-specific negative density dependence (Bagchi *et al*., 2010, 2014; Liu *et al*., 2012, i.e., fungicide increases seedling survival at high conspecific densities). Extending these experiments to include habitat variation would directly test the JC–HP interaction by asking whether natural enemies suppress seedlings most strongly in habitats where they would otherwise have high survival. In practice, this predicts a positive correlation between the increase in survival due to fungicide (Δ*S*_fungicide_) and baseline survival in the absence of enemies (*S*_baseline_). Equivalently, this corresponds to Cov(Δ*S*_fungicide_, *S*_baseline_) *>* 0, directly paralleling the JC–HP covariance term.

### Future directions and conclusion

Several simplifying assumptions in the model suggest avenues for extension. I considered a one-dimensional habitat axis, but real forests contain multiple abiotic gradients that vary independently and across scales (John *et al*., 2007). Spatiotemporal, rather than static, abiotic variation is also common tropical forests (Wright, 2002). Similarly, JC-effect strength was modeled as independent of habitat type, although evidence suggests that habitat variables can modify the baseline strength of density dependence (e.g. Yang *et al*., 2023). Certain environments (e.g., moister habitats; Milici *et al*., 2020) may therefore generate stronger JC–HP interactions. JC-effects may also persist long after adult death (Magee *et al*., 2024, 2025), which could modify JC-HP interactions. I also assumed natural enemies to be perfectly specialized, whereas spillover across related species is common (Gilbert & Webb, 2007; Ødegaard *et al*., 2005). Incorporating partial enemy overlap may alter JC–HP interactions depending on whether species that share habitat preferences also share natural enemies (see Stump, 2017). The coevolution of natural enemy specialization on host trees and tree habitat specialization (Fine *et al*., 2004, 2013) is a particularly interesting avenue to explore in future models. Finally, habitat partitioning is implemented through a single pre-adult recruitment/establishment component (i.e., all habitat dependence enters before individuals enter the adult population), whereas in real forests habitat can also affect adult demographic rates; allowing habitat dependence at the adult stage could alter adult spatial distributions and thus where JC-effects are generated, although the qualitative role of spatial alignment between habitat suitability and density-dependent suppression should persist.

Functional traits such as seed size trade off with susceptibility to density dependence: larger-seeded species experience weaker conspecific negative density dependence (Lebrija-Trejos *et al*., 2016). This axis of variation could be integrated into the tolerance–fecundity trade-off (see Supplementary Information; Muller-Landau, 2010; D’Andrea & O’Dwyer, 2021). Interspecific variation in dispersal distances (Muller-Landau *et al*., 2008) provides another avenue to link JC–effects (and their interactions with habitat partitioning) to other life-history trade-offs (Stump & Comita, 2020). Together, these axes of variation could help embed JC–HP interactions into a broader trait-based framework.

While these extensions may yield further insights, the main result is likely robust: JC-effects and habitat partitioning are not independent forces but interact strongly through the spatial structure of forests. When habitats are spatially autocorrelated, habitat specialization generates conspecific aggregation in favorable environments, concentrating JC-effects where they matter most and producing a powerful mechanism for coexistence. By contrast, aggregation arising from dispersal limitation undermines coexistence. Thus, the consequences of aggregation depend critically on its underlying cause. Recognizing this distinction highlights a clear path forward. Identifying and quantifying the processes that generate spatial patterns, and integrating these with process-based theory, will be essential for linking local dynamics to the maintenance of community-level biodiversity. The results presented here offer a tractable basis to connect spatial data from forest plots and experiments with theoretical predictions, providing a concrete framework to test how local processes scale to sustain tropical forest diversity.

## Data availability

All code required to reproduce the analyses and results in this paper is available at: https://github.com/DanielSmithEcology/Janzen_Connell_Habitat_Partitioning_Interactions.

## Acknowledgements

This research was funded by the U.S. National Science Foundation grant NSF OCE 1851489 and National Science Foundation Division of Environmental Biology Award 2240430. I thank J. Timothy Wootton, Catherine A Pfister, Mercedes Pascual, Trevor Price, Lukas Magee, and the Wootton-Pfister lab for insightful discussion and feedback on the manuscript.

