## Supplementary Information for "Janzen-Connell effects and habitat-induced aggregation synergistically promote species coexistence"

### Supplemental information for: Janzen-Connell effects and habitat-induced aggregation synergistically promote species coexistence

4 Interspecific variation in baseline JC-effect strength 27

5 Confirmation of dynamic equilibrium 29

6 Analysis of approximations 30

7 Parameter tables 34

### Introduction

This supplement is divided into several sections. First, I provide a more detailed derivation of the model presented in the main text. Second, I derive the approximate invasion criterion from the main text and present the associated derivations. Third, I provide details and results for the tolerance–fecundity trade-off model referenced in the main text. Fourth, I present supplementary simulations in which species exhibit interspecific variation in baseline JC-effect strength, $a$. Fifth, I provide numerical evidence (visualizations) showing that the simulations presented in the main text reach a dynamic equilibrium. Sixth, I present numerical results showing that the approximations depicted in the main text are, to a reasonable degree, accurate. Seventh, I present two parameter tables showing variables, parameters, and symbols that appear in the main text and those that appear only in the Supplementary Information, respectively.

### 1 General model derivation and details

To connect intuition with the compact model presented in the main text, I begin with a mechanistic description of offspring dynamics and its numerical implementation. The community consists of a grid of $P = X \times X$ patches, each occupied by a single adult individual. Species are indexed by locally $i = 1, \ldots, N$ where $N$ is the total number in the local pool; a total of $L$ species reside in the regional pool ($L \geq N$). At each time step, adults produce propagules that are dispersed across patches. These propagules then experience mortality from two independent sources: habitat filtering and conspecific negative density dependence (i.e., JC-effects), in addition to baseline density-independent mortality. Because the numerical simulations allow dispersal to be either global or dispersal-limited, I also provide a detailed description below of how local dispersal kernels and dispersal limitation are implemented computationally.

#### 1.1 Dispersal

Let $d_i(x)$ denote the *dispersal pressure* of species $i$ at patch $x$: the kernel-weighted contribution of adults of species $i$ to patch $x$, independent of fecundity. If each adult of species $i$ produces $f_i$ propagules per time step, then the expected number (density) of propagules of species $i$ arriving at patch $x$ immediately after dispersal is

$$f_i \, d_i(x).$$

(Baseline density-independent survival is applied after dispersal; for compactness below I later combine $f_i$ and baseline survival into an intrinsic fitness parameter $Y_i$.)

Dispersal can be either global or local. In the global case, propagules are distributed uniformly across all patches. If species $i$ occupies a fraction $p_i$ of patches, dispersal pressure is spatially constant:

$$d_i(x) \;=\; p_i, \tag{1}$$

so that the expected arriving propagule density is $f_i\, d_i(x) = p_i f_i$.

With local dispersal, dispersal pressure is determined by a discrete kernel defined over Chebyshev distance. For two patches with coordinates $(a_1, b_1)$ and $(a_2, b_2)$, the Chebyshev distance is

$$z \;=\; \max\big(|a_1 - a_2|,\, |b_1 - b_2|\big).$$

To construct the discrete kernel, I begin from an isotropic two-dimensional Gaussian dispersal kernel and use its radial displacement distribution (a Rayleigh form). Let $\zeta$ denote Euclidean distance between patch centers. The radial probability density is

$$k(\zeta; \sigma_D) \;=\; \frac{2\zeta}{\sigma_D^2} \exp\!\left(-\frac{\zeta^2}{\sigma_D^2}\right), \tag{2}$$

where $\sigma_D$ is the dispersal scale (in grid-cell units) shared across species. I then bin this continuous kernel into unit-width distance shells and convert it into a per-cell probability for each Chebyshev shell. Specifically, defining $W(z)$ as the probability that a propagule lands in a particular patch at Chebyshev distance $z$ from its source patch, I set

$$W(0) \;=\; \int_0^{1/2} k(\zeta; \sigma_D)\, d\zeta, \qquad W(z) \;=\; \frac{1}{8z} \int_{z-1/2}^{z+1/2} k(\zeta; \sigma_D)\, d\zeta \quad (z \geq 1), \tag{3}$$

so that the total probability mass assigned to shell $z \geq 1$ is $8z\, W(z)$, distributed uniformly across the $8z$ patches in that shell.

Let $\mathbf{1}\{s(y) = i\}$ indicate whether patch $y$ is occupied by species $i$. Under local dispersal, dispersal pressure at patch $x$ is

$$d_i(x) \;=\; \sum_{y \in \text{grid}} \mathbf{1}\{s(y) = i\}\, W\big(\text{dist}(x, y)\big), \tag{4}$$

where $\text{dist}(x, y)$ denotes Chebyshev distance between patches $x$ and $y$. The expected arriving propagule density is then $f_i\, d_i(x)$.

In practice, I truncate the kernel at a finite maximum distance to reduce computational cost and to approximate rare long-distance dispersal. Let $z_{\max}$ denote the largest Chebyshev shell computed explicitly (set by a user-specified distance cap and/or by requiring that the cumulative mass captured by the discrete kernel exceed a high threshold). Defining the captured (local) mass as

$$D_{\text{Lim}} \;=\; W(0) + \sum_{z=1}^{z_{\max}} 8z\, W(z),$$

the remaining tail mass $1 - D_{\text{Lim}}$ is distributed uniformly across the grid. Under this approximation,

$$d_i(x) \;\approx\; \sum_{y \in \text{grid}} \mathbf{1}\{s(y) = i\}\, W\big(\text{dist}(x,y)\big) \;+\; (1 - D_{\text{Lim}})\, p_i, \tag{5}$$

so that the expected arriving propagule density is $f_i\, d_i(x)$ while still allowing rare long-distance arrivals at low computational cost.

##### 1.2 Mortality dynamics

Let $S_i(x,t)$ be the number of surviving propagules of species $i$ on patch $x$ at time $t$. These propagules experience two independent sources of mortality:

$$\frac{dS_i(x,t)}{dt} = -\ell_i(x)\, S_i(x,t) \;-\; j_i(x,t)\, S_i(x,t), \tag{6}$$

where $\ell_i(x)$ is the density-independent mortality rate due to baseline survival and abiotic habitat mismatch and $j_i(x,t)$ is the density-dependent mortality rate from JC-effects. The initial condition is $S_i(x,0) = d_i(x) f_i$.

Solving over the vulnerable period of length $\tau$ gives

$$S_i(x,\tau) = d_i(x) f_i\, \exp\big(-\ell_i(x)\tau\big)\, \exp\big(-j_i(x)\tau\big). \tag{7}$$

For compactness, I write $S_i(x)$ for $S_i(x,\tau)$.

##### 1.3 Habitat effects

Density-independent mortality *rate* is modeled as a quadratic function of the local habitat value $E(x) \in [0,1]$:

$$\ell_i(x)\tau = m_i + \frac{(E(x) - h_{\text{opt},i})^2}{2\sigma_h^2}, \tag{8}$$

for which mortality rate is lowest at $h_{\text{opt}}$. Then,

$$\exp\big(-\ell_i(x)\tau\big) = e^{-m_i}\, \exp\left(-\frac{(E(x) - h_{\text{opt},i})^2}{2\sigma_h^2}\right).$$

The first factor, $e^{-m_i}$, is baseline survival on the optimal habitat, while the second defines a Gaussian response curve. Define

$$H_i(x) = \exp\left(-\frac{(E(x) - h_{\text{opt},i})^2}{2\sigma_h^2}\right). \tag{9}$$

I also consider the stress–tolerance fecundity trade-off, which is discussed in the following section.

##### 1.4 JC-effects

Let $A_i(M(x))$ be the number of conspecific adults of species $i$ in the $M \times M$ Moore neighborhood of patch $x$. JC-effects impose mortality by adding linearly to the *per-patch cumulative hazard* over the vulnerable period,

$$j_i(x)\tau = a_i A_i(M(x)), \tag{10}$$

so that survival from JC-effects on patch $x$ is

$$\exp\big(-j_i(x)\tau\big) = \exp\big(-a_i A_i(M(x))\big). \tag{11}$$

I define

$$J_i(x) = 1 - \exp\big(-a_i A_i(M(x))\big), \tag{12}$$

so that $1 - J_i(x) = \exp\big(-a_i A_i(M(x))\big)$ is the survival probability from JC-effects. For intuition, when a single conspecific adult is present ($A_i = 1$), the per-patch mortality probability is $1 - e^{-a_i}$. I use the term "JC-effect (predation) pressure" to denote the *spatially summed* cumulative hazard contributed by a single adult across its $M \times M$ footprint, which is $a_i M^2$.

##### 1.5 Overall seedling survival

Combining all factors, I obtain

$$S_i(x) = d_i(x)\, f_i e^{-m_i}\, H_i(x)\,(1 - J_i(x)). \tag{13}$$

Finally, I define intrinsic fitness as

$$Y_i = f_i e^{-m_i}, \tag{14}$$

so that the survival function takes the compact form used in the main text:

$$S_i(x) = d_i(x)\, Y_i\, H_i(x)\,(1 - J_i(x)). \tag{15}$$

##### 1.6 Expression for population growth

For any patch-level quantity $g(x)$, define the spatial average

$$\mathbb{E}_x[g(x)] := \frac{1}{P}\sum_{x=1}^{P} g(x). \tag{16}$$

Then, putting all the above different pieces together, the population dynamics of species $i$ are described by

$$p_i(t+1) = p_i(t)(1-\delta) + \delta\,\mathbb{E}_x\left[\frac{S_i(x)}{\sum_j S_j(x)}\right], \tag{17}$$

where $p_i(t)$ is the proportion of patches occupied by species $i$ at time step $t$, and $\delta$ is the adult mortality probability.

### 2 Invasion criteria

#### 2.1 JC-effects and habitat partitioning, no dispersal limitation

I consider invasion by a rare focal species $i$ with habitat optimum $h_{\text{opt},i} \in [0,1]$ into a resident community of $N$ species indexed by $j = 1,\ldots,N$. Throughout this subsection, the invader index $i$ is not in $\{1,\ldots,N\}$. Let $p_j(t)$ denote resident frequencies for $j = 1,\ldots,N$, and let $p_i(t)$ denote the invader frequency, with $\sum_{j=1}^{N} p_j(t) + p_i(t) = 1$. For compactness, I use $\sum_{\text{all } j}$ to denote a sum over the $N$ residents *and* the invader (i.e., $j \in \{1,\ldots,N\} \cup \{i\}$). Let $\lambda_i = p_i(t+1)/p_i(t)$ be the finite rate of increase of the invader.

$$\lambda_i = (1-\delta) + \delta\,\mathbb{E}_x\left[\frac{Y_i\,H_i(x)\big(1-J_i(x)\big)}{\sum_{\text{all } j} p_j\,Y_j\,H_j(x)\big(1-J_j(x)\big)}\right]. \tag{18}$$

I consider the weighted growth rate of species $i$, $\tilde{\lambda}_i = (\lambda_i - 1)/\delta > 0$. This is given by

$$\tilde{\lambda}_i = \mathbb{E}_x\left[\frac{Y_i\,H_i(x)\big(1-J_i(x)\big)}{\sum_{\text{all } j} p_j\,Y_j\,H_j(x)\big(1-J_j(x)\big)}\right] - 1, \tag{19}$$

where $Y_i$ is intrinsic fitness, $H_i(x)$ is the habitat response at location $x$, $J_i(x)$ is the Janzen–Connell (JC) effect at $x$, and the expectation is taken over all space $x$. Write $\mu_{H_i(x)} = \mathbb{E}_x[H_i(x)]$ (I keep species-specific $\mu_{H_i(x)}$ for now; the identical-$\mu_{H(x)}$ case is noted later). Note that I have written the sum as an expectation over $x$, $\mathbb{E}_x$, which is more convenient notation here.

**Step 1: Normalize by $\mu_{H_i(x)}$, $Y_i$, and $(1 - J_i(x))$.** I divide the numerator and denominator inside the expectation by $Y_i(1 - J_i(x))\,\mu_{H_i(x)}$:

$$\tilde{\lambda}_i = \mathbb{E}_x\left[\frac{H_i(x)}{\mu_{H_i(x)}}\,\frac{1}{\sum_{\text{all } j} p_j\,\frac{Y_j}{Y_i}\,\frac{H_j(x)}{\mu_{H_i(x)}}\,\frac{1-J_j(x)}{1-J_i(x)}}\right] - 1. \tag{20}$$

This operation allows several terms to cancel. I define the inner sum as

$$I(x) \;:=\; \sum_{\text{all } j} p_j \, \frac{Y_j}{Y_i} \, \frac{H_j(x)}{\mu_{H_i(x)}} \, \frac{1 - J_j(x)}{1 - J_i(x)}. \tag{21}$$

Then the reciprocal factor is $F(x) = 1/I(x)$, and (20) becomes

$$\tilde{\lambda}_i = \mathbb{E}_x\left[\frac{H_i(x)}{\mu_{H_i(x)}} \, F(x)\right] - 1, \tag{22}$$

which allows for easier simplifications.

**Step 2: First-order expansion of $F(x) = 1/I(x)$ around $I(x) = 1$.** I expand $g(s) = 1/s$ at $s = 1$: $g(s) \approx g(1) + g'(1)(s-1) = 1 - (s-1) = 2 - s$. Hence,

$$F(x) \;=\; \frac{1}{I(x)} \;\approx\; 2 - I(x), \tag{23}$$

which I use below in the approximations.

**Step 3: Log-linearize $I(x)$.** I write each multiplicative factor in (21) as an exponential and linearize at the neutral point $Y_j \approx Y_i$, $H_j(x) \approx \mu_{H_i(x)}$, and $J_j(x) \approx J_i(x)$:

$$\frac{Y_j}{Y_i} \, \frac{H_j(x)}{\mu_{H_i(x)}} \, \frac{1 - J_j(x)}{1 - J_i(x)} \;=\; \exp\Big( \ln Y_j - \ln Y_i + \ln \tfrac{H_j(x)}{\mu_{H_i(x)}} + \ln(1 - J_j(x)) - \ln(1 - J_i(x)) \Big). \tag{24}$$

Using $\exp(u) \approx 1 + u$ for small $u$,

$$\frac{Y_j}{Y_i} \, \frac{H_j(x)}{\mu_{H_i(x)}} \, \frac{1 - J_j(x)}{1 - J_i(x)} \;\approx\; 1 + \big(\ln Y_j - \ln Y_i\big) + \ln \tfrac{H_j(x)}{\mu_{H_i(x)}} + \big(\ln(1 - J_j(x)) - \ln(1 - J_i(x))\big). \tag{25}$$

Averaging over species with weights $p_j$ (and noting that $\sum_{\text{all } j} p_j = 1$) gives

$$\begin{aligned} I(x) &= \sum_{\text{all } j} p_j \Big[ 1 + (\ln Y_j - \ln Y_i) + \ln \tfrac{H_j(x)}{\mu_{H_i(x)}} + \ln(1 - J_j(x)) - \ln(1 - J_i(x)) \Big] \\ &= 1 + \Big( \overline{\ln Y_j}^{\,p_j} - \ln Y_i \Big) + \sum_{\text{all } j} p_j \ln \tfrac{H_j(x)}{\mu_{H_i(x)}} + \Big( \overline{\ln(1 - J_j(x))}^{\,p_j} - \ln(1 - J_i(x)) \Big), \end{aligned} \tag{26}$$

where e.g. $\overline{\ln Y_j}^{\,p_j} = \sum_{\text{all } j} \ln Y_j \, p_j$ and $\overline{\ln(1 - J_j(x))}^{\,p_j} = \sum_{\text{all } j} \ln(1 - J_j(x)) \, p_j$ (i.e., proportion-scaled quantities). These quanities will cancel out later.

**Step 4: Small-variation approximations for habitat effects and JC-effects.** Starting from (26),

$$I(x) = 1 + \left(\overline{\ln Y_j}^{p_j} - \ln Y_i\right) + \sum_{\text{all } j} p_j \ln \frac{H_j(x)}{\mu_{H_i(x)}} + \left(\overline{\ln(1 - J_j(x))}^{p_j} - \ln(1 - J_i(x))\right), \tag{27}$$

I apply the first-order approximations $\ln z \approx z - 1$ (for $z$ near 1) and $\ln(1-u) \approx -u$ (for small $u$):

$$\sum_{\text{all } j} p_j \ln \frac{H_j(x)}{\mu_{H_i(x)}} \;\approx\; \sum_{\text{all } j} p_j \left(\frac{H_j(x)}{\mu_{H_i(x)}} - 1\right) \;=\; \sum_{\text{all } j} p_j \frac{H_j(x)}{\mu_{H_i(x)}} - 1, \tag{28}$$

and

$$\overline{\ln(1 - J_j(x))}^{p_j} - \ln(1 - J_i(x)) \;\approx\; -\overline{J_j(x)}^{p_j} + J_i(x). \tag{29}$$

Therefore,

$$I(x) \;\approx\; \left(\overline{\ln Y_j}^{p_j} - \ln Y_i\right) \;+\; \sum_{\text{all } j} p_j \frac{H_j(x)}{\mu_{H_i(x)}} \;-\; \overline{J_j(x)}^{p_j} \;+\; J_i(x), \tag{30}$$

where the $+1$ and $-1$ terms have canceled.

**Step 5: Substitute $F(x)$ with $2 - I(x)$ and collect terms.** With $F(x) = 1/I(x) \approx 2 - I(x)$, substituting terms yields

$$\tilde{\lambda}_i = \mathbb{E}_x\left[\frac{H_i(x)}{\mu_{H_i(x)}}\, F(x)\right] - 1 \;\approx\; 1 - \mathbb{E}_x\left[\frac{H_i(x)}{\mu_{H_i(x)}}\, I(x)\right]. \tag{31}$$

From (30),

$$I(x) \;\approx\; \left(\overline{\ln Y_j}^{p_j} - \ln Y_i\right) \;+\; \sum_{\text{all } j} p_j \frac{H_j(x)}{\mu_{H_i(x)}} \;-\; \overline{J_j(x)}^{p_j} \;+\; J_i(x), \tag{32}$$

so

$$\begin{aligned}
\tilde{\lambda}_i \approx{}& 1 - \mathbb{E}_x\left[\frac{H_i(x)}{\mu_{H_i(x)}}\left(\overline{\ln Y_j}^{p_j} - \ln Y_i\right)\right] - \mathbb{E}_x\left[\frac{H_i(x)}{\mu_{H_i(x)}} \sum_{\text{all } j} p_j \frac{H_j(x)}{\mu_{H_i(x)}}\right] + \mathbb{E}_x\left[\frac{H_i(x)}{\mu_{H_i(x)}}\, \overline{J_j(x)}^{p_j}\right] \\
&- \mathbb{E}_x\left[\frac{H_i(x)}{\mu_{H_i(x)}} J_i(x)\right] \\
={}& \quad 1 + \ln Y_i - \overline{\ln Y_j}^{p_j} - \mathbb{E}_x\left[\frac{H_i(x)}{\mu_{H_i(x)}} \sum_{\text{all } j} p_j \frac{H_j(x)}{\mu_{H_i(x)}}\right] + \mathbb{E}_x\left[\frac{H_i(x)}{\mu_{H_i(x)}}\, \overline{J_j(x)}^{p_j}\right] - \mathbb{E}_x\left[\frac{H_i(x)}{\mu_{H_i(x)}} J_i(x)\right],
\end{aligned} \tag{33}$$

where $\mathbb{E}_x[H_i(x)/\mu_{H_i(x)}] = 1$ is used to pull out the $p_j$-weighted means in the final expression.

**Step 6: Equilibrium cleanup.** From this point onward, I evaluate expressions at the resident equilibrium (i.e., in the rare-invader limit $p_i \to 0$), so that $\sum_{\text{all } j}$ reduces to $\sum_{j=1}^{N}$ and the $p_j$ are resident frequencies with $\sum_{j=1}^{N} p_j = 1$. At equilibrium, the residents have zero growth. Because the invader index $i$ is not among the residents ($i \notin \{1, \ldots, N\}$), resident averages are written as $\frac{1}{N}\sum_{k=1}^{N}(\cdot)$. Thus, as noted by Chesson (1994),

$$\tilde{\lambda}_i \;-\; \frac{1}{N}\sum_{k=1}^{N}\tilde{\lambda}_k \;=\; \tilde{\lambda}_i. \tag{34}$$

From (33), the invader's first–order growth is

$$\tilde{\lambda}_i \;\approx\; \big(\ln Y_i - \overline{\ln Y}^{p_j}\big) - \mathbb{E}_x\!\left(\frac{H_i(x)}{\mu_{H_i(x)}}\sum_{j=1}^{N} p_j \frac{H_j(x)}{\mu_{H_i(x)}}\right) + \mathbb{E}_x\!\left(\frac{H_i(x)}{\mu_{H_i(x)}}\,\overline{J(x)}^{p_j}\right) - \mathbb{E}_x\!\left(\frac{H_i(x)}{\mu_{H_i(x)}} J_i(x)\right), \tag{35}$$

and, for a resident $k$,

$$\tilde{\lambda}_k \;\approx\; \big(\ln Y_k - \overline{\ln Y}^{p_j}\big) - \mathbb{E}_x\!\left(\frac{H_k(x)}{\mu_{H_k(x)}}\sum_{j=1}^{N} p_j \frac{H_j(x)}{\mu_{H_k(x)}}\right) + \mathbb{E}_x\!\left(\frac{H_k(x)}{\mu_{H_k(x)}}\,\overline{J(x)}^{p_j}\right) - \mathbb{E}_x\!\left(\frac{H_k(x)}{\mu_{H_k(x)}} J_k(x)\right). \tag{36}$$

Averaging (36) over $k = 1, \ldots, N$ and subtracting from (35) gives

$$\begin{aligned}
\tilde{\lambda}_i \approx\; & \underbrace{\Big[\big(\ln Y_i - \overline{\ln Y}^{p_j}\big) - \big(\tfrac{1}{N}\sum_{k=1}^{N}\ln Y_k - \overline{\ln Y}^{p_j}\big)\Big]}_{=\;\ln Y_i \;-\; \frac{1}{N}\sum_{k=1}^{N}\ln Y_k} \\
& - \left[\mathbb{E}_x\!\left(\frac{H_i(x)}{\mu_{H_i(x)}}\sum_{j=1}^{N} p_j \frac{H_j(x)}{\mu_{H_i(x)}}\right) - \tfrac{1}{N}\sum_{k=1}^{N}\mathbb{E}_x\!\left(\frac{H_k(x)}{\mu_{H_k(x)}}\sum_{j=1}^{N} p_j \frac{H_j(x)}{\mu_{H_k(x)}}\right)\right] \\
& + \left[\mathbb{E}_x\!\left(\frac{H_i(x)}{\mu_{H_i(x)}}\,\overline{J(x)}^{p_j}\right) - \tfrac{1}{N}\sum_{k=1}^{N}\mathbb{E}_x\!\left(\frac{H_k(x)}{\mu_{H_k(x)}}\,\overline{J(x)}^{p_j}\right)\right] \\
& - \left[\mathbb{E}_x\!\left(\frac{H_i(x)}{\mu_{H_i(x)}} J_i(x)\right) - \tfrac{1}{N}\sum_{k=1}^{N}\mathbb{E}_x\!\left(\frac{H_k(x)}{\mu_{H_k(x)}} J_k(x)\right)\right].
\end{aligned} \tag{37}$$

*JC mean–coupling bracket cancels under no association.* For any species $s$,

$$\mathbb{E}_x\!\left(\frac{H_s(x)}{\mu_{H_s(x)}}\,\overline{J(x)}^{p_j}\right) = \underbrace{\mathbb{E}_x\!\left(\frac{H_s(x)}{\mu_{H_s(x)}}\right)}_{=1}\mathbb{E}_x[\overline{J(x)}^{p_j}] \;+\; \mathrm{Cov}_x\!\left(\frac{H_s(x)}{\mu_{H_s(x)}}, \overline{J(x)}^{p_j}\right). \tag{38}$$

Hence,

$$\begin{aligned}&\mathbb{E}_x\Big(\tfrac{H_i(x)}{\mu_{H_i(x)}}\,\overline{J(x)}^{p_j}\Big) - \tfrac{1}{N}\sum_{k=1}^{N}\mathbb{E}_x\Big(\tfrac{H_k(x)}{\mu_{H_k(x)}}\,\overline{J(x)}^{p_j}\Big) = \\ &\underbrace{\Big[1 - \tfrac{1}{N}\sum_{k=1}^{N} 1\Big]\,\mathbb{E}_x[\overline{J(x)}^{p_j}]}_{=0} + \Big[\mathrm{Cov}_x\big(\tfrac{H_i(x)}{\mu_{H_i(x)}}, \overline{J(x)}^{p_j}\big) - \tfrac{1}{N}\sum_{k=1}^{N}\mathrm{Cov}_x\big(\tfrac{H_k(x)}{\mu_{H_k(x)}}, \overline{J(x)}^{p_j}\big)\Big].\end{aligned} \tag{39}$$

If habitat use is uncorrelated with the spatial mean JC-effect for all species, i.e., $\mathrm{Cov}_x\big(\frac{H_s(x)}{\mu_{H_s(x)}}, \overline{J(x)}^{p_j}\big) = 0$, then the entire bracket in the second line of (37) is

$$\mathbb{E}_x\Big(\tfrac{H_i(x)}{\mu_{H_i(x)}}\,\overline{J(x)}^{p_j}\Big) - \tfrac{1}{N}\sum_{k=1}^{N}\mathbb{E}_x\Big(\tfrac{H_k(x)}{\mu_{H_k(x)}}\,\overline{J(x)}^{p_j}\Big) = 0. \tag{40}$$

This assumption is reasonable, although rare scenarios could allow for exceptions.

Combining everything from (40) and (34), (37) reduces to

$$\begin{aligned}\tilde{\lambda}_i = \ &\Big[\ln Y_i \ - \ \tfrac{1}{N}\sum_{k=1}^{N}\ln Y_k\Big] \\ &- \left[\mathbb{E}_x\left(\tfrac{H_i(x)}{\mu_{H_i(x)}}\sum_{j=1}^{N} p_j\tfrac{H_j(x)}{\mu_{H_i(x)}}\right) - \tfrac{1}{N}\sum_{k=1}^{N}\mathbb{E}_x\left(\tfrac{H_k(x)}{\mu_{H_k(x)}}\sum_{j=1}^{N} p_j\tfrac{H_j(x)}{\mu_{H_k(x)}}\right)\right] \\ &- \left[\mathbb{E}_x\Big(\tfrac{H_i(x)}{\mu_{H_i(x)}}J_i(x)\Big) - \tfrac{1}{N}\sum_{k=1}^{N}\mathbb{E}_x\Big(\tfrac{H_k(x)}{\mu_{H_k(x)}}J_k(x)\Big)\right].\end{aligned} \tag{41}$$

which will simplify when I take the limit $p_i \to 0$ below.

**Step 7: Define $\overline{h(x)}$ and decompose the resident JC term.** Motivated by the habitat bracket in (41), define the post-contrast habitat adjustment

$$\overline{h(x)} \ = \ \frac{1}{N}\sum_{k=1}^{N}\mathbb{E}_x\left[\frac{H_k(x)}{\mu_{H_k(x)}}\sum_{\text{all } j} p_j\frac{H_j(x)}{\mu_{H_k(x)}}\right] \ - \ \mathbb{E}_x\left[\frac{H_i(x)}{\mu_{H_i(x)}}\sum_{\text{all } j} p_j\frac{H_j(x)}{\mu_{H_i(x)}}\right]. \tag{42}$$

This will be simplified in the below section, but it describes the direct impact of habitats partitioning on invasion.

Invader rarity, $p_i \to 0$, implies

$$\mathbb{E}_x\left(\frac{H_i(x)}{\mu_{H_i(x)}}J_i(x)\right) = 0, \tag{43}$$

because, under global dispersal, species $i$ does not experience JC-effects. Thus, equation (41) becomes

$$\tilde{\lambda}_i \;\approx\; \Big[\ln Y_i \;-\; \tfrac{1}{N}\sum_{k=1}^{N}\ln Y_k\Big] \;+\; \overline{h(x)} \;+\; \tfrac{1}{N}\sum_{k=1}^{N}\mathbb{E}_x\left(\frac{H_k(x)}{\mu_{H_k(x)}}J_k(x)\right). \tag{44}$$

For the final term, I use the property $\mathbb{E}_x[AB] = \mathbb{E}_x[A]\cdot\mathbb{E}_x[B] + \mathrm{Cov}_x(A,B)$. Then, for each resident $k$,

$$\mathbb{E}_x\left(\frac{H_k(x)}{\mu_{H_k(x)}}J_k(x)\right) = \underbrace{\mathbb{E}_x\big[J_k(x)\big]}_{\text{spatial mean JC}} \;+\; \underbrace{\mathrm{Cov}_x\left(J_k(x),\,\frac{H_k(x)}{\mu_{H_k(x)}}\right)}_{\text{JC–HP covariance}}, \tag{45}$$

because $\mathbb{E}_x[H_k/\mu_{H_k(x)}] = 1$. Averaging (45) over residents $k = 1,\dots,N$ yields

$$\tfrac{1}{N}\sum_{k=1}^{N}\mathbb{E}_x\left(\tfrac{H_k}{\mu_{H_k(x)}}J_k\right) = \overline{\mathbb{E}_x[J(x)]} \;+\; \overline{\mathrm{Cov}_x}\left(J(x),\,\frac{H(x)}{\mu_{H(x)}}\right), \tag{46}$$

where the bar denotes the mean over residents.

**Final invasion condition.** Substituting (46) into (44) gives

$$\boxed{\tilde{\lambda}_i \;\approx\; \ln Y_i \;-\; \overline{\ln Y} \;+\; \overline{h(x)} \;+\; \overline{\mathbb{E}_x[J(x)]} \;+\; \overline{\mathrm{Cov}_x}\left(J(x),\,\frac{H(x)}{\mu_{H(x)}}\right)} \tag{47}$$

noting $\overline{\ln Y} = \frac{1}{N}\sum_{k=1}^{N}\ln Y_k$. This is the expression from the main text. Below (after the derivation of the dispersal-limitation case), I derive simplified expressions for $\overline{h(x)}$, $\overline{\mathbb{E}_x[J(x)]}$, and the upper bound of $\overline{\mathrm{Cov}_x}\big(J(x),\,H(x)/\mu_{H(x)}\big)$.

#### 2.2 JC-effects, dispersal limitation, and no habitat partitioning

I now consider the case where species exhibit dispersal limitation but no habitat partitioning. This initially takes the form

$$\tilde{\lambda}_i = \mathbb{E}_x\left[\frac{Y_i\,d_i(x)\big(1-J_i(x)\big)}{\sum_j p_j Y_j\,d_j(x)\big(1-J_j(x)\big)}\right] - 1. \tag{48}$$

However, for analytical tractability, I consider an approximate model.

First, I simplify the form of $d_i(x)$. Let $d$ denote the proportion of propagules that escape JC-effects within the $M \times M$ Moore neighborhood generated by their parent tree. Then, it suffices to distinguish between locally dispersed and non-locally dispersed propagules.

Second, I introduce what I call the "pre-dispersal effect" model. Rather than tracking locally dispersed propagules *per se*, I assume that a fraction $1-d$ of each parent's propagules experiences heightened mortality before dispersal, and that all propagules are subsequently dispersed uniformly over space. I use the notation $J_i^L(x)$ to denote the additional reduction in survival experienced by propagules harmed by the parent tree. $J_i(x)$ simply refers to propagules that were *not* harmed by their parent tree. This yields the simplified expression:

$$\tilde{\lambda}_i \approx (1-d)\,\mathbb{E}_x\left[\frac{Y_i\big(1-J_i^L(x)\big)}{\sum_j S_j(x)}\right] + d\cdot\mathbb{E}_x\left[\frac{Y_i\big(1-J_i(x)\big)}{\sum_j S_j(x)}\right] - 1, \tag{49}$$

where $\sum_j S_j(x)$ is the total abundance of seeds arriving at patch $x$.

Next, I apply the approximation $\ln(A/B) \approx 1 + \ln A - \ln B$, rewriting each term inside the expectations:

$$\begin{aligned}\tilde{\lambda}_i \approx (1-d)\,\mathbb{E}_x&\left[1+\ln Y_i + \ln\big(1-J_i^L(x)\big) - \ln\sum_j S_j(x)\right]\\ &+ d\cdot\mathbb{E}_x\left[1+\ln Y_i + \ln\big(1-J_i(x)\big) - \ln\sum_j S_j(x)\right] - 1.\end{aligned} \tag{50}$$

Using $\ln(1-A) \approx -A$, this simplifies to

$$\tilde{\lambda}_i \approx (1-d)\,\mathbb{E}_x\left[1+\ln Y_i - J_i^L(x) - \ln\sum_j S_j(x)\right] + d\cdot\mathbb{E}_x\left[1+\ln Y_i - J_i(x) - \ln\sum_j S_j(x)\right] - 1. \tag{51}$$

Collecting terms gives

$$\tilde{\lambda}_i \approx \ln Y_i - \mathbb{E}_x\big[(1-d)J_i^L(x) + d\,J_i(x)\big] - \mathbb{E}_x\left[\ln\overline{S(x)}\right], \tag{52}$$

where $\ln\overline{S(x)}$ denotes the mean log total seed input.

As in the above derivation, I use the fact that at equilibrium the finite rate of increase of each of the $N$ resident species is zero, so I again use the property

$$\tilde{\lambda}_i - \tfrac{1}{N}\sum_{k\neq i}\tilde{\lambda}_k \;=\; \tilde{\lambda}_i. \tag{53}$$

See Chesson (1994). This yields

$$\begin{aligned}\tilde{\lambda}_i \approx\; &\ln Y_i - \tfrac{1}{N}\sum_{k\neq i}\ln Y_k + \tfrac{1}{N}\sum_{k\neq i}\mathbb{E}_x\big[(1-d)J_k^L(x) + d\,J_k(x)\big] - \mathbb{E}_x\big[(1-d)J_i^L(x) + d\,J_i(x)\big]\\ &+ \tfrac{1}{N}\sum_{k\neq i}\mathbb{E}_x\left[\ln\overline{S(x)}\right] - \mathbb{E}_x\left[\ln\overline{S(x)}\right].\end{aligned} \tag{54}$$

The last two terms cancel. Taking $p_i \to 0$ (so that $\mathbb{E}_x[J_i(x)] \to 0$, since only non-local effects remain) gives

$$\tilde{\lambda}_i \approx \ln Y_i - \overline{\ln Y} + (1-d)\,\overline{\mathbb{E}_x[J^L(x)]} - d\,\overline{\mathbb{E}_x[J(x)]} - (1-d)\,\mathbb{E}_x[J_i^L(x)], \tag{55}$$

where overbars denote averages across species.

Finally, note that local JC-effects are equivalent to adding one additional conspecific in the Moore neighborhood. Let $J^L(x)' = 1 - J^L(x)$ and $J(x)' = 1 - J(x)$ denote survival probabilities. Then

$$J^L(x)' = J(x)' e^{-a},$$

and for the rare species,

$$J_i^L(x)' = e^{-a},$$

since $J_i(x)' = 1$ in the non-local case. Substituting back, $J_i^L(x) = 1 - e^{-a}$, the probability of mortality due to local JC-effects. Thus,

$$\tilde{\lambda}_i \approx \ln Y_i - \overline{\ln Y} + (1-d)\,\overline{\mathbb{E}_x[1 - J(x)'e^{-a}]} - d\,\overline{\mathbb{E}_x[1 - J(x)']} - (1-d)(1-e^{-a}). \tag{56}$$

After some algebraic simplifications,

$$\boxed{\tilde{\lambda}_i \approx \ln Y_i - \overline{\ln Y} + \overline{\mathbb{E}_x[J(x)]}\Big(d + (1-d)e^{-a}\Big)} \tag{57}$$

which is equation (14) from the main text.

#### 2.3 Evaluation of $\overline{\mathbb{E}_x[J(x)]}$

Here, I show the derivations of $\overline{\mathbb{E}_x[J(x)]}$ from the main text. Recall that the goal here is a tractable approximation, not an exact expression. Therefore, I make several linearizations and simplifications below.

##### 2.3.1 Evaluation of $\overline{\mathbb{E}_x[J(x)]}$ assuming random mixing

I evaluate the resident-mean JC effect by first computing $\mathbb{E}[J_j(x)]$ for a given species $j$ under random mixing (with no explicit spatial autocorrelation).

Consider a focal patch $x$ and its $M \times M$ Moore neighborhood. Define

$$A_j(x) \;=\; \sum_{m \in \mathcal{N}_M(x)} \mathbf{1}\{\text{the adult at site } m \text{ is species } j\}, \tag{58}$$

so that $A_j(x)$ is the count of conspecific adults around $x$. JC imposes a survival *penalty* that increases with $A_j(x)$:

$$J_j(x) \;=\; 1 - \exp\bigl(-\,a\,A_j(x)\bigr), \tag{59}$$

so the survival factor entering the demographic kernel is $1 - J_j(x) = \exp(-a\,A_j(x))$. Here $a > 0$ controls JC strength.

Under random mixing with species-$j$ occupancy $p_j$ per site and independence across sites,

$$A_j(x) \sim \mathrm{Bin}\bigl(M^2,\; p_j\bigr). \tag{60}$$

**Expectation $\overline{\mathbb{E}_x[J_j(x)]}$.** Using (59) and (60),

$$\begin{aligned}
\mathbb{E}\bigl[J_j(x)\bigr] &= \mathbb{E}\bigl[\,1 - e^{-aA_j(x)}\bigr] = 1 - \mathbb{E}\bigl[e^{-aA_j(x)}\bigr] \\
&= 1 - \sum_{y=0}^{M^2} \binom{M^2}{y} (p_j)^y (1-p_j)^{M^2-y} e^{-ay} \\
&= 1 - \bigl((1-p_j) + p_j e^{-a}\bigr)^{M^2} \\
&= 1 - \bigl(1 - (1-e^{-a})\,p_j\bigr)^{M^2}.
\end{aligned} \tag{61}$$

Equation (61) is *exact* for the binomial model.

**Small-parameter linearization.** From (61),

$$\mathbb{E}\bigl[J_j(x)\bigr] \;\approx\; 1 - \exp\bigl(-\,(1-e^{-a})\,M^2 p_j\bigr).$$

When $(1-e^{-a})\,M^2 p_j$ is small, this can be linearized via

$$\exp(-z) \approx 1 - z \quad\Rightarrow\quad \mathbb{E}\bigl[J_j(x)\bigr] \;\approx\; (1-e^{-a})\,M^2\,p_j. \tag{62}$$

**Resident mean.** Let $N$ be the number of residents (excluding the rare invader), and take a simple average over residents:

$$\overline{\mathbb{E}_x[J(x)]} \;:=\; \frac{1}{N}\sum_{j\neq i} \mathbb{E}_x\bigl[J_j(x)\bigr] \;\approx\; \frac{1}{N}\sum_{j\neq i}(1-e^{-a})\,M^2\,p_j.$$

Because the invader is rare, $\sum_{j\neq i} p_j \to 1$, so

$$\boxed{\overline{\mathbb{E}_x[J(x)]} \;\approx\; (1-e^{-a})\,\frac{M^2}{N}} \tag{63}$$

which is the same as equation (7) from the main text.

##### 2.3.2 Moment-based approximation of $\overline{\mathbb{E}_x[J(x)]}$

The random-mixing result above assumes $A_j(x) \sim \mathrm{Bin}(M^2, p_j)$. To avoid this assumption, I approximate $\mathbb{E}[J_j(x)]$ directly from the first two moments of the $M \times M$ conspecific count:

$$A_j(x) = \sum_{m \in \mathcal{N}_M(x)} \mathbf{1}\{\text{adult at } m \text{ is } j\}, \qquad J_j(x) = 1 - e^{-aA_j(x)}.$$

Let

$$\mu_{j,M} := \mathbb{E}[A_j], \qquad \sigma^2_{j,M} := \mathrm{Var}(A_j), \qquad \kappa_{j,M} := \frac{\sigma^2_{j,M}}{\mu_{j,M}},$$

so that $\kappa_{j,M}$ is the local variance-to-mean ratio (aggregation index). This formulation places no constraint on the spatial process that generates $A_j$; it only requires that $\mu_{j,M}$ and $\sigma^2_{j,M}$ exist.

**Second-order approximation.** For $f(y) = e^{-ay}$, a second-order Taylor expansion about $y = \mu_{j,M}$ gives

$$\mathbb{E}\big[e^{-aA_j}\big] \approx e^{-a\mu_{j,M}} \left(1 + \tfrac{1}{2}a^2\sigma^2_{j,M}\right).$$

Therefore,

$$\mathbb{E}[J_j(x)] = 1 - \mathbb{E}\big[e^{-aA_j}\big] \approx 1 - e^{-a\mu_{j,M}}\Big(1 + \tfrac{1}{2}a^2\sigma^2_{j,M}\Big) = 1 - e^{-a\mu_{j,M}}\Big(1 + \tfrac{1}{2}a^2\,\mu_{j,M}\,\kappa_{j,M}\Big). \tag{64}$$

**Small-$a$ series linearization** Taking the linearization of the exponential term in (64) yields:

$$\mathbb{E}[J_j(x)] \approx a\,\mu_{j,M} - \tfrac{1}{2}a^2\left(\mu^2_{j,M} + \mu_{j,M}\,\kappa_{j,M}\right). \tag{65}$$

Next, I substitute the mean proportions into the equation 65. $\mu_{j,M} = M^2 p_j$. Writing the result in terms of $p_j$ and $\kappa_{j,M}$:

$$\mathbb{E}[J_j(x)] \approx aM^2 p_j - \tfrac{1}{2}a^2\left(M^4 p_j^2 + M^2 p_j\,\kappa_{j,M}\right). \tag{66}$$

**Resident mean.** Averaging over residents ($N$ species excluding the rare invader) gives with the above expansion yields:

$$\overline{\mathbb{E}_x[J(x)]} \approx \underbrace{aM^2 \frac{1}{N}\sum_{j \neq i} p_j}_{aM^2/N} - \frac{a^2}{2}\left[M^4 \frac{1}{N}\sum_{j \neq i} p_j^2 + M^2 \frac{1}{N}\sum_{j \neq i} p_j\,\kappa_{j,M}\right]. \tag{67}$$

When residents are near the average abundance ($p_j \approx 1/N$), $\frac{1}{N}\sum p_j^2 \approx 1/N^2$ and $\frac{1}{N}\sum p_j\,\kappa_{j,M} \approx \frac{1}{N}\overline{\kappa}_M$, where

$$\overline{\kappa}_M := \frac{1}{N}\sum_{k\neq i}\kappa_{k,M}$$

is the community mean aggregation at scale $M$. Then

$$\overline{\mathbb{E}_x[J(x)]} \;\approx\; \frac{aM^2}{N} \;-\; \frac{a^2}{2}\left(\frac{M^4}{N^2} \;+\; \frac{M^2}{N}\,\overline{\kappa}_M\right). \tag{68}$$

When $N$ is large and $M^2 \ll N$ (noting this must be true for the above linearization to hold) the $\frac{M^4}{N^2}$ term is negligible, giving

$$\boxed{\overline{\mathbb{E}_x[J(x)]} \;\approx\; \frac{aM^2}{N}\left(1-\tfrac{a}{2}\,\overline{\kappa}_M\right)} \tag{69}$$

which is the same as equation (8) from the main text.

##### 2.4 Derivation of $\overline{h(x)}$

I derive the expressions for $\overline{h(x)}$ used in the main text. Let $H_s(x)$ denote the (Gaussian) habitat response of species $s$ evaluated at location $x$. As before

$$\mu_{H_s(x)} \;=\; \mathbb{E}_x[H_s(x)].$$

There are $N$ residents plus a rare invader $i$. Recall that the residents are indexed by $k = 1, \ldots, N$ (so $i \notin \{1, \ldots, N\}$). I use $\sum_{\text{all } j}$ to denote a sum over the $N$ residents *and* the invader (i.e., $j \in \{1, \ldots, N\} \cup \{i\}$), with

$$\sum_{\text{all } j} p_j = 1, \qquad \text{and in the invasion limit } p_i \to 0,\ \sum_{j=1}^{N} p_j = 1.$$

**Definition.** I begin from

$$\overline{h(x)} \;:=\; \underbrace{\frac{1}{N}\sum_{k=1}^{N}\mathbb{E}_x\left[\frac{H_k(x)}{\mu_{H_k(x)}}\sum_{\text{all } j} p_j\frac{H_j(x)}{\mu_{H_k(x)}}\right]}_{\text{Term B}} \;-\; \underbrace{\mathbb{E}_x\left[\frac{H_i(x)}{\mu_{H_i(x)}}\sum_{\text{all } j} p_j\frac{H_j(x)}{\mu_{H_i(x)}}\right]}_{\text{Term A}}. \tag{70}$$

**Step 1: Covariance decomposition (no random–optima assumption).** For each pair of species $(s, j)$, define

$$\mu_{H_s(x)} = \mathbb{E}_x[H_s(x)], \qquad \sigma^2_{H_s(x)} = \mathrm{Var}_x\big(H_s(x)\big), \qquad C_{sj} = \mathrm{Cov}_x\big(H_s(x), H_j(x)\big). \tag{71}$$

Then

$$\mathbb{E}_x\big[H_s(x)H_j(x)\big] = \mu_{H_s(x)}\mu_{H_j(x)} + C_{sj}. \tag{72}$$

**Step 2: Terms A and B in covariance form (denominators consistent with** (70)**).** Because (70) normalizes the inner sum by the *focal* mean (i.e., by $\mu_{H_i(x)}$ in Term A and by $\mu_{H_k(x)}$ in Term B), the relevant identity is

$$\frac{\mathbb{E}_x[H_sH_j]}{\mu^2_{H_s(x)}} = \frac{\mu_{H_j(x)}}{\mu_{H_s(x)}} + \frac{C_{sj}}{\mu^2_{H_s(x)}}. \tag{73}$$

Using (73),

$$\text{Term A} = \sum_{\text{all } j} p_j \, \frac{\mathbb{E}_x[H_iH_j]}{\mu^2_{H_i(x)}} = \sum_{\text{all } j} p_j \left[\frac{\mu_{H_j(x)}}{\mu_{H_i(x)}} + \frac{C_{ij}}{\mu^2_{H_i(x)}}\right].$$

Separating the $j = i$ contribution,

$$\text{Term A} = \sum_{j=1}^{N} p_j \left[\frac{\mu_{H_j(x)}}{\mu_{H_i(x)}} + \frac{C_{ij}}{\mu^2_{H_i(x)}}\right] + p_i \left[1 + \frac{C_{ii}}{\mu^2_{H_i(x)}}\right],$$

so in the invasion limit $p_i \to 0$ the final term is negligible.

Similarly,

$$\text{Term B} = \frac{1}{N}\sum_{k=1}^{N}\sum_{\text{all } j} p_j \, \frac{\mathbb{E}_x[H_kH_j]}{\mu^2_{H_k(x)}} = \frac{1}{N}\sum_{k=1}^{N}\sum_{\text{all } j} p_j \left[\frac{\mu_{H_j(x)}}{\mu_{H_k(x)}} + \frac{C_{kj}}{\mu^2_{H_k(x)}}\right].$$

Again, the $j = i$ contribution inside the inner sum is $O(p_i)$ and vanishes as $p_i \to 0$.

**Step 3: Combine Terms A and B.** Taking the invasion limit $p_i \to 0$ (so $\sum_{\text{all } j} \to \sum_{j=1}^{N}$) yields

$$\overline{h(x)} = \frac{1}{N}\sum_{k=1}^{N}\sum_{j=1}^{N} p_j \left[\frac{\mu_{H_j(x)}}{\mu_{H_k(x)}} + \frac{C_{kj}}{\mu^2_{H_k(x)}}\right] - \sum_{j=1}^{N} p_j \left[\frac{\mu_{H_j(x)}}{\mu_{H_i(x)}} + \frac{C_{ij}}{\mu^2_{H_i(x)}}\right]. \tag{74}$$

Under the common–mean/exchangeability approximation used below (i.e., $\mu_{H_s(x)} \approx \mu_{H(x)}$ for all species including the invader), the mean–ratio terms satisfy $\mu_{H_j(x)}/\mu_{H_k(x)} \approx 1$ and $\mu_{H_j(x)}/\mu_{H_i(x)} \approx 1$, so the constant parts cancel and

$$\overline{h(x)} = \frac{1}{N}\sum_{k=1}^{N}\sum_{j=1}^{N} p_j \, \frac{C_{kj}}{\mu^2_{H(x)}} - \sum_{j=1}^{N} p_j \, \frac{C_{ij}}{\mu^2_{H(x)}}. \tag{75}$$

This is the form used for the symmetric simplifications in Steps 4–5.

**Step 4: Symmetric/exchangeable simplification.** Assume residents are exchangeable with common mean $\mu_{H(x)}$ and variance $\sigma^2_{H(x)}$, and define an average correlation $\bar{\rho}$ so that

$$C_{kk} = \sigma^2_{H(x)}, \qquad \overline{C_{kj}} = \bar{\rho}\,\sigma^2_{H(x)} \quad (k \neq j),$$

where the bar indicates an average over partners. Because residents are exchangeable, the particular weights $\{p_j\}$ no longer matter (only that they sum to one). Substituting gives

$$\frac{1}{N}\sum_{k=1}^{N} p_k \frac{C_{kk}}{\mu^2_{H(x)}} = \frac{1}{N}\frac{\sigma^2_{H(x)}}{\mu^2_{H(x)}},$$

$$\frac{1}{N}\sum_{k=1}^{N}\sum_{j\neq k} p_j \frac{C_{kj}}{\mu^2_{H(x)}} = \frac{N-1}{N}\,\bar{\rho}\,\frac{\sigma^2_{H(x)}}{\mu^2_{H(x)}},$$

$$\sum_{j=1}^{N} p_j \frac{C_{ij}}{\mu^2_{H(x)}} = \bar{\rho}\,\frac{\sigma^2_{H(x)}}{\mu^2_{H(x)}}.$$

Hence

$$\overline{h(x)} = \left(\frac{\sigma^2_{H(x)}}{\mu^2_{H(x)}}\right)\left[\frac{1}{N} + \frac{N-1}{N}\,\bar{\rho} \;-\; \bar{\rho}\right]. \tag{76}$$

This simplifies to the result

$$\boxed{\overline{h(x)} = \frac{1}{N}\left(\frac{\sigma^2_{H(x)}}{\mu^2_{H(x)}}\right)(1-\bar{\rho}).} \tag{77}$$

as shown in the main text.

Furthermore, independent responses across species ($\bar{\rho} = 0$) yield

$$\overline{h(x)} = \frac{1}{N}\frac{\sigma^2_{H(x)}}{\mu^2_{H(x)}},$$

while increasing average alignment ($\bar{\rho} \to 1$) suppresses $\overline{h(x)}$ toward zero.

**Step 5: Random/independent optima & Gaussian closed form.** As noted above, when species' optima are independently and uniformly distributed, cross–covariances vanish on average, so $\bar{\rho} = 0$. For the Gaussian response, I further assume $\sigma_h$ is sufficiently small that nearly the entire density of $H(x)$ falls within $x \in [0, 1]$, so edge effects can be neglected and the integrals can be evaluated as on an infinite domain.

$$H(x) = \exp\Big(-\frac{(x-h)^2}{2\sigma_h^2}\Big) \text{ on a large domain, } \mu_{H(x)} = \int H(x)\,dx = \sigma_h\sqrt{2\pi}, \quad \mathbb{E}_x[H(x)^2] = \sigma_h\sqrt{\pi}.$$

Therefore

$$\frac{\sigma^2_{H(x)}}{\mu^2_{H(x)}} = \frac{\mathbb{E}[H^2] - \mu^2_{H(x)}}{\mu^2_{H(x)}} = \frac{\mathbb{E}[H^2]}{\mu^2_{H(x)}} - 1 = \frac{\sigma_h\sqrt{\pi}}{(\sigma_h\sqrt{2\pi})^2} - 1 = \frac{1}{2\sqrt{\pi}\,\sigma_h} - 1.$$

Plugging into (77) with $\bar{\rho} = 0$ gives

$$\boxed{\overline{h(x)} = \frac{1}{N}\left(\frac{1}{2\sqrt{\pi}\,\sigma_h} - 1\right).} \tag{78}$$

Equivalently, since $\mu_{H(x)} = \sigma_h\sqrt{2\pi}$,

$$\frac{1}{2\sqrt{\pi}\,\sigma_h} = \frac{1}{\sqrt{2}\,\mu_{H(x)}}, \qquad \Rightarrow \qquad \overline{h(x)} = \frac{1}{N}\left(\frac{1}{\sqrt{2}\,\mu_{H(x)}} - 1\right),$$

which matches the main text expressions.

##### 2.4.1 Evenly spaced optima (regular niche packing)

In the main text I note that $\overline{h(x)}$ can decline more rapidly with community size $N$ when resident habitat optima are *regularly spaced*, i.e. approximately evenly distributed along the habitat axis with adjacent optima separated by $\sim 1/N$.

To obtain a fully explicit expression, I use a top-hat ("box") approximation to the Gaussian habitat response on a periodic habitat axis $x \in [0, 1)$:

$$H_j(x) = \mathbf{1}\Big\{d_{\text{per}}(x, h_{\text{opt},j}) \le \frac{w}{2}\Big\}, \qquad h_{\text{opt},j} = \frac{j-1}{N}, \quad j = 1, \ldots, N, \tag{79}$$

where $d_{\text{per}}(x, h_{\text{opt}}) := \min\{|x - h_{\text{opt}}|,\ 1 - |x - h_{\text{opt}}|\}$ is the periodic distance and $w$ preserves the Gaussian area,

$$w = \int_{-\infty}^{\infty} \exp\Big(-\frac{(x - h_{\text{opt},i})^2}{2\sigma_h^2}\Big)\,dx = \sigma_h\sqrt{2\pi}. \tag{80}$$

For $x$ uniform on $[0, 1)$, define $\mu_{H_j(x)} := \mathbb{E}_x[H_j(x)]$. Under the edge-free (periodic) approximation, $\mu_{H_j(x)} = w$ for all $j$ (on a bounded non-periodic domain, $\mu_{H_j(x)} \le w$ for optima near the boundary). In what follows I assume the edge-free case and write the common mean as $\mu_{H(x)} = w$. I also assume $w \le 1/2$, so each box occupies at most half of the periodic habitat axis.

Assume a symmetric resident community, $p_j = 1/N$. Using the same definition of $\overline{h(x)}$ as above (Term B – Term A),

$$\overline{h(x)} = \frac{1}{N^2 \mu_{H(x)}^2} \sum_{k=1}^{N} \sum_{j=1}^{N} \mathbb{E}_x\big[H_k(x) H_j(x)\big] \; - \; 1, \tag{81}$$

where the $-1$ arises because for a "typical" invader (averaging over a uniformly placed invader optimum $h_{\text{opt},i} \sim U[0,1)$, independent of the resident lattice) one has $\mathbb{E}_{x,h_{\text{opt},i}}[H_i(x)H_j(x)] = \mu_{H(x)}^2$ for any resident $j$, hence Term A equals 1.

For the box kernel, $\mathbb{E}_x[H_k(x)H_j(x)]$ is the length of overlap of two intervals of length $\mu_{H(x)}$ centered at $h_k$ and $h_j$. Writing $d_{kj} = d_{\text{per}}(h_k, h_j)$ for the periodic distance between optima (and using $w \le 1/2$),

$$\mathbb{E}_x\big[H_k(x) H_j(x)\big] = \max\big(0,\ \mu_{H(x)} - d_{kj}\big)\,. \tag{82}$$

With evenly spaced optima, pairwise distances are $d = m/N$ for integers $m$. Let

$$\alpha := \mu_{H(x)} N, \qquad q := \lfloor \alpha \rfloor, \qquad r := \alpha - q \in [0,1), \tag{83}$$

so that $q$ is the largest integer spacing index with positive overlap (i.e. the number of neighbor spacings that fit within width $\mu_{H(x)}$) and $r$ is the remainder. Evaluating (81) gives

$$\overline{h(x)} = \frac{r(1-r)}{\mu_{H(x)}^2 N^2}. \tag{84}$$

This is nonnegative and oscillatory in $N$ through $r = \{\mu_{H(x)} N\}$, with

$$0 \le \overline{h(x)} \le \frac{1}{4\,\mu_{H(x)}^2 N^2} \qquad \text{(maximum at } r = \tfrac{1}{2}\text{; zero whenever } \mu_{H(x)} N \in \mathbb{Z}). \tag{85}$$

However, the oscillatory nature of this approximation of $\overline{h(x)}$ is likely an artifact of the top-hat approximation. To obtain a smooth representative value, I average over the "phase" $r$ (which cycles through $[0,1)$ as $N$ varies, or equivalently if $\mu_{H(x)}$ varies slightly across realizations), so that $r$ is effectively uniform on $[0,1)$. Since $\mathbb{E}[r(1-r)] = 1/6$, this yields

$$\mathbb{E}\big[\overline{h(x)}\big] = \frac{1}{6\,\mu_{H(x)}^2 N^2} = \frac{1}{6\,w^2 N^2}. \tag{86}$$

Thus, under regular spacing, $\overline{h(x)}$ can decay as $O(N^{-2})$, in contrast to the $O(N^{-1})$ scaling obtained under randomly distributed optima.

##### 2.5 Upper bound on JC–HP covariance

In this section, I derive an upper bound for the JC–HP covariance term. The derivation assumes equal specialization $\mu_{H_i(x)} = \mu_{H_k(x)} = \mu_{H(x)}$ and approximately equal species abundances, although the intuition applies more broadly.

The goal is to bound

$$\overline{\mathrm{Cov}_x}\Big(J(x), \tfrac{H(x)}{\mu_{H(x)}}\Big) \;=\; \frac{1}{N}\sum_{k=1}^{N} \mathrm{Cov}_x\Big(J_k(x), \tfrac{H_k(x)}{\mu_{H_k(x)}}\Big), \qquad \mu_{H_k(x)} \equiv \mu_{H(x)}, \tag{87}$$

using the Cauchy–Schwarz inequality. For each species,

$$\Big|\mathrm{Cov}_x\Big(J_k(x), \tfrac{H_k(x)}{\mu_{H(x)}}\Big)\Big| \;\le\; \sigma_{J_k(x)}\,\sigma_{H_k(x)/\mu_{H(x)}}, \tag{88}$$

where $\sigma$ denotes the spatial standard deviation. Since $\sigma_{H(x)/\mu_{H(x)}} = \sigma_{H(x)}/\mu_{H(x)}$, this implies

$$\Big|\overline{\mathrm{Cov}_x}\Big(J(x), \tfrac{H(x)}{\mu_{H(x)}}\Big)\Big| \;\le\; \frac{\sigma_{J(x)}\,\sigma_{H(x)}}{\mu_{H(x)}}. \tag{89}$$

In what follows, I derive an upper bound for $\sigma_{J(x)}$ and an explicit expression for $\sigma_{H(x)}$.

###### 2.5.1 Heuristic upper bound on JC–HP covariance

I first derive a simple, scaling-transparent bound that I will show before the tighter bound.

Let $A_i(x)$ be the number of conspecific adults of species $i$ in the $M \times M$ neighborhood at location $x$, with

$$\mu_{i,M} \;=\; \mathbb{E}_x[A_i(x)], \qquad \kappa_{i,M} \;=\; \frac{\mathrm{Var}_x\big(A_i(x)\big)}{\mu_{i,M}} \quad \text{(mean–variance index of aggregation).}$$

The JC mortality is $J_i(x) = 1 - e^{-aA_i(x)}$. Using that $J'(A) = ae^{-aA} \le a$ (Lipschitz with constant $a$) and a delta–method linearization,

$$\sigma_{J_i(x)} \;\lesssim\; a\,\sigma_{A_i(x)} \;=\; a\,\sqrt{\kappa_{i,M}\,\mu_{i,M}}.$$

Averaging over species and applying Cauchy–Schwarz to $\frac{1}{N}\sum_i \sqrt{\kappa_{i,M}\mu_{i,M}}$ gives

$$\overline{\sigma_{J(x)}} \;=\; \frac{1}{N}\sum_{i=1}^{N} \sigma_{J_i(x)} \;\lesssim\; a\,\sqrt{\,\overline{\kappa}_M\,\overline{\mu}_M\,}, \qquad \overline{\kappa}_M = \frac{1}{N}\sum_i \kappa_{i,M}, \;\; \overline{\mu}_M = \frac{1}{N}\sum_i \mu_{i,M}.$$

325 Under approximately equal abundances, $\mu_{i,M} \approx M^2/N$ so $\overline{\mu}_M \approx M^2/N$. Hence

$$\overline{\sigma_{J(x)}} \lesssim a \frac{M}{\sqrt{N}} \sqrt{\overline{\kappa}_M}.$$

326 Now apply Cauchy–Schwarz to the spatial covariance and normalize habitat by its spatial 327 mean:

$$\left|\overline{\mathrm{Cov}_x}\Big(J(x), \tfrac{H(x)}{\mu_{H(x)}}\Big)\right| \leq \overline{\sigma_{J(x)}}\, \sigma_{H(x)/\mu_{H(x)}} = \overline{\sigma_{J(x)}} \frac{\sigma_{H(x)}}{\mu_{H(x)}}.$$

328 Combining the two expressions yields the heuristic bound

$$\boxed{\overline{\mathrm{Cov}_x}\Big(J(x), \tfrac{H(x)}{\mu_{H(x)}}\Big) \lesssim \underbrace{\tfrac{aM}{\sqrt{N}}\sqrt{\overline{\kappa}_M}}_{\approx \sigma_{J(x)}} \times \underbrace{\tfrac{\sigma_{H(x)}}{\mu_{H(x)}}}_{\mathrm{CV}(H(x))}}. \tag{90}$$

329 This is equation (12) in the main text. Thus, stabilization from JC–HP covariance increases with 330 mean intraspecific aggregation $\overline{\kappa}_M$ and with the coefficient of variation of habitat effects, while 331 decreasing with regional richness $N$. This contrasts with the purely "direct" JC effect, whose 332 strength declines with aggregation.

##### 333 2.5.2 Tighter JC-HP upper bound

334 Below, I derive equation (13) from the main text.

335 **Upper bound on the variance of $J(x)$** The key term in (89) is $\sigma_{J(x)}$. Recall that here $J(x)$ 336 denotes the *mortality probability* due to JC-effects:

$$J(x) = 1 - e^{-aA(x)},$$

337 where $A(x)$ is the number of conspecific adults in an $M \times M$ Moore neighborhood. Since 338 $A(x) \in [0, M^2]$ and $\mathbb{E}[A(x)] = M^2/N$, on average each species contributes $M^2/N$ adults to the 339 neighborhood.

340 Because the exponential is convex, the variance of $J(x)$ is maximized when $A(x)$ takes only 341 extreme values. Specifically, consider

$$A(x) = \begin{cases} 0 & \text{with prob. } \frac{N-1}{N}, \\ M & \text{with prob. } \frac{1}{N}, \end{cases}$$

which satisfies the mean constraint. Writing $q = e^{-aM}$, the induced distribution of $J(x)$ is

$$J(x) = \begin{cases} 0 & \text{with prob. } \frac{N-1}{N}, \\ 1-q & \text{with prob. } \frac{1}{N}. \end{cases}$$

**Expectation.**

$$\mathbb{E}[J(x)] = \frac{N-1}{N} \cdot 0 + \frac{1}{N} \cdot (1-q) = \frac{1-q}{N}.$$

**Second moment.**

$$\mathbb{E}[J(x)^2] = \frac{N-1}{N} \cdot 0^2 + \frac{1}{N} \cdot (1-q)^2 = \frac{(1-q)^2}{N}.$$

**Variance.** Subtracting gives

$$\begin{aligned} \mathrm{Var}[J(x)] &= \mathbb{E}[J(x)^2] - \big(\mathbb{E}[J(x)]\big)^2 && (91) \\ &= \frac{(1-q)^2}{N} - \left(\frac{1-q}{N}\right)^2 && (92) \\ &= (1-q)^2 \left(\frac{1}{N} - \frac{1}{N^2}\right). && (93) \end{aligned}$$

Therefore,

$$\sigma_{J(x)} \;\le\; (1 - e^{-aM^2}) \sqrt{\tfrac{1}{N} - \tfrac{1}{N^2}}. \tag{94}$$

This bound gives the maximum spatial variability in JC-induced mortality.

**Approximate variance of $H(x)$.** Now consider the habitat response

$$H(x) = \exp\!\left(-\frac{(x-\mu)^2}{2\sigma_h^2}\right), \qquad X \sim \mathrm{Unif}[0,1], \qquad \mu_{H(x)} = \sigma_h\sqrt{2\pi}. \tag{95}$$

When $\sigma_h \ll 1$ and $\mu$ lies well within the habitat range, the Gaussian bump around $\mu$ is narrow compared to the unit interval. Thus the uniform average over $[0,1]$ can be well-approximated by extending the integral to the whole real line, with negligible boundary error. In this regime the required moments reduce to Gaussian integrals:

$$\mathbb{E}[H(X)] \;\approx\; \int_{-\infty}^{\infty} e^{-(x-\mu)^2/(2\sigma_h^2)}\, dx = \sigma_h\sqrt{2\pi} \;=\; \mu_{H(x)},$$

and

$$\mathbb{E}[H(X)^2] \;\approx\; \int_{-\infty}^{\infty} e^{-(x-\mu)^2/\sigma_h^2}\, dx = \sigma_h\sqrt{\pi}.$$

**Variance and standard deviation (direct form).** Subtracting $\mu_{H(x)}^2$ gives

$$\mathrm{Var}[H(X)] \;\approx\; \sqrt{\pi}\,\sigma_h - 2\pi\sigma_h^2 \;=\; \frac{\mu_{H(x)}}{\sqrt{2}} - \mu_{H(x)}^2. \tag{96}$$

Hence

$$\sigma_{H(x)} \;\approx\; \sqrt{\tfrac{\mu_{H(x)}}{\sqrt{2}} - \mu_{H(x)}^2}, \qquad \sigma_{H(x)/\mu_{H(x)}} \;\approx\; \sqrt{\tfrac{1}{\mu_{H(x)}\sqrt{2}} - 1}. \tag{97}$$

**Variance in CV form.** Equivalently, writing in terms of the coefficient of variation:

$$\frac{\sigma_{H(x)}^2}{\mu_{H(x)}^2} = \frac{\mathbb{E}[H^2]}{\mu_{H(x)}^2} - 1 = \frac{1}{\sqrt{2}\,\mu_{H(x)}} - 1. \tag{98}$$

Thus

$$\mathrm{CV}_{H(x)}^2 \;\approx\; \frac{1}{\sqrt{2}\,\mu_{H(x)}} - 1. \tag{99}$$

**Final upper bound** Substituting the bounds for $\sigma_{J(x)}$ and $\sigma_{H(x)/\mu_{H(x)}}$ into (89) gives

$$\overline{\mathrm{Cov}_x}\Big(J(x), \tfrac{H(x)}{\mu_{H(x)}}\Big) \;\lesssim\; (1 - e^{-aM^2})\,\sqrt{\tfrac{1}{N}\left(1 - \tfrac{1}{N}\right)}\,\sqrt{\tfrac{1}{\mu_{H(x)}\sqrt{2}} - 1} \tag{100}$$

which is the expression in equation (13) of the main text. This inequality provides an explicit upper bound on the magnitude of JC–HP covariance. It holds under the assumptions of narrow niche widths ($\mu_{H(x)} \ll 1$), approximately equal species abundances, and identical $\mu_{H(x)}$ across species.

#### 3 Tolerance–fecundity trade-off model

Empirical work suggests that small-seeded species tend to be more fecund but less tolerant of stressful environments, whereas large-seeded species are less fecund but more stress tolerant (Moles *et al.*, 2003; Moles & Westoby, 2004; Muller-Landau, 2010; D'Andrea *et al.*, 2013; D'Andrea & O'Dwyer, 2021). To connect this pattern to the framework presented in the main text (and above), I consider a specialist–generalist trade-off in which species differ jointly in fecundity and environmental tolerance ("stress tolerance").

##### 3.1 Model specifications

Let $x \in [0,1]$ denote a one-dimensional habitat (stress) gradient. Species are indexed by their fecundity rank $R_i \in \{1, \ldots, S\}$, with $R_i = 1$ the most fecund species and $R_i = S$ the least

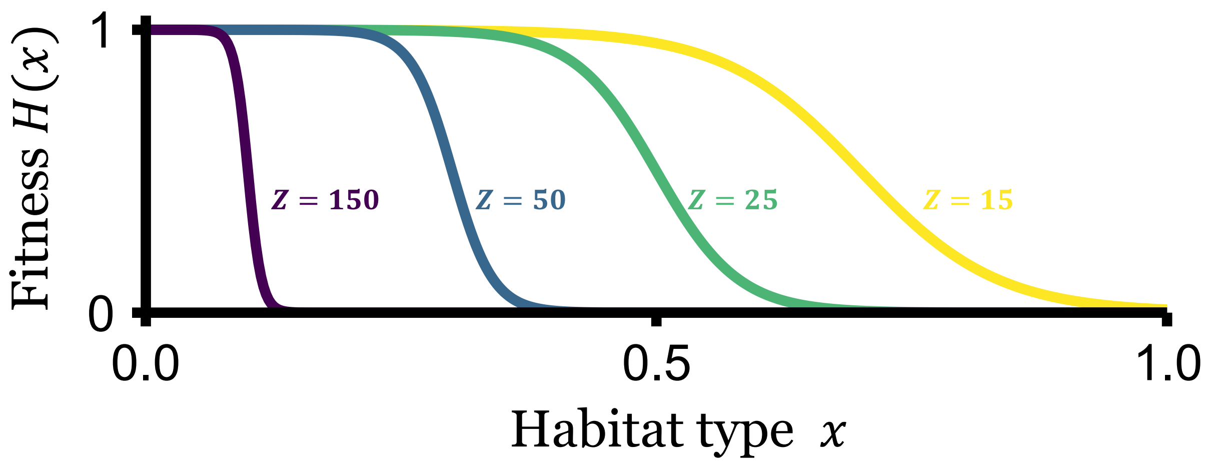

Figure 1: **Tolerance–fecundity trade-off generates Z-shaped habitat responses.** Curves show the habitat survival function $H_i(x) = 1/\Big(1 + \exp\big(-Z[b_0 + R_i/S - x]\big)\Big)$ for species with different fecundity–tolerance combinations. Four values of $R_i/S = \{0.1,\ 0.3,\ 0.5,\ 0.7\}$ under four steepness parameters $Z = \{150, 50, 25, 15\}$, respectively. The baseline threshold was set to $b_0 = 0.0$, so the point of steep decline occurs at $x \approx R_i/S$. Higher $R_i/S$ (higher fecundity) produces narrower tolerance, while lower $R_i/S$ produces broader tolerance. The habitat axis spans $x \in [0, 1]$, and $H_i(x)$ ranges between 0 (no survival) and 1 (full survival).

fecund (e.g., obtained by sorting lognormal fecundity draws). The probability that a propagule of species $i$ establishes on a patch of type $x$ is

$$H_i(x) \;=\; \frac{1}{1 + \exp\Big(-Z\big[b_0 + \frac{R_i}{S} - x\big]\Big)}. \tag{101}$$

Here $b_0 \in [0, 1]$ is a baseline tolerance intercept (the fraction of the gradient tolerated by a mid-ranked species when $Z$ is large), and $Z > 0$ controls the steepness of the transition from low to high tolerance along the gradient. Larger $R_i$ (lower fecundity) shifts the logistic to the right, so less fecund species tolerate more stressful (larger-$x$) environments. See Fig. 1 for a visualization.

Equation (101) produces "Z-shaped" tolerance curves across species ranks: (i) as $Z \to \infty$, $H_i(x) \to \mathbf{1}\{x < b_0 + R_i/S\}$, recovering a step-function tolerance limit akin to Muller-Landau (2010); (ii) as $Z \to 0$, $H_i(x) \to 1/2$ for all $i$ and all $x$, eliminating habitat filtering; (iii) increasing $b_0$ shifts all species to tolerate a larger fraction of the gradient.

Thus, the trade-off is encoded by a monotone fecundity ordering (equalizing differences) coupled with a monotone tolerance shift (stabilizing via habitat partitioning).

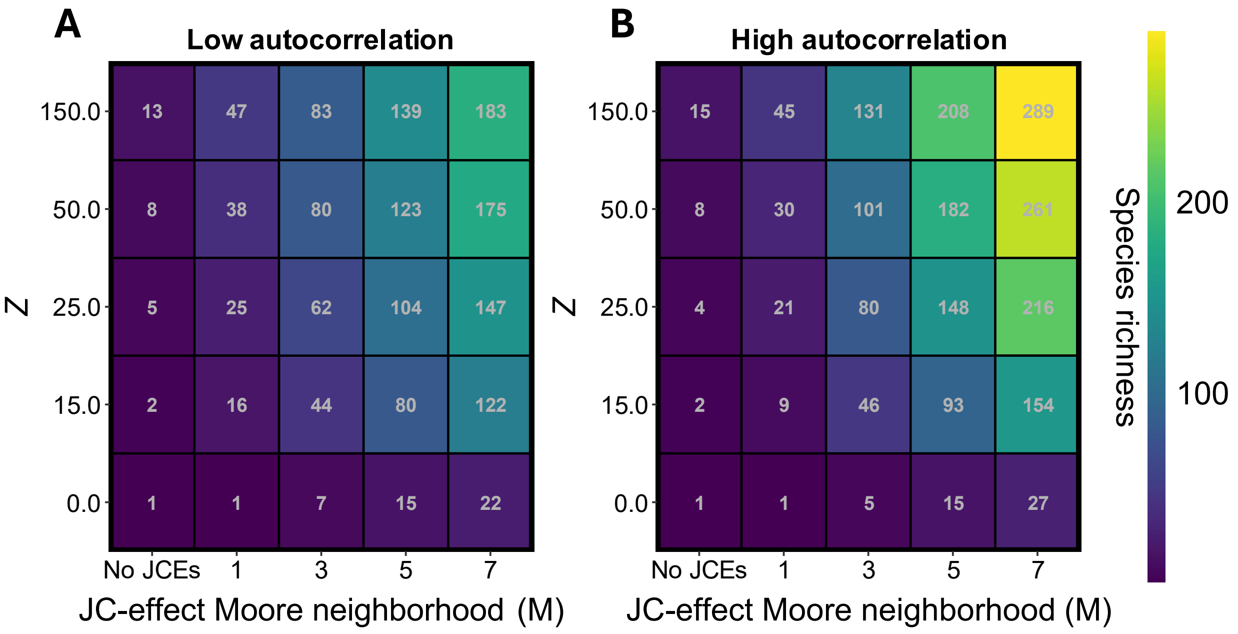

Figure 2: **Positive habitat spatial autocorrelation promotes higher species richness when JC-effects operate alongside the tolerance–fecundity trade-off.** Plots show equilibrium species richness (color scale) across combinations of JC-effect scale ($M$, x-axis) and niche breadth ($\sigma_h$, y-axis). (A) Low spatial autocorrelation (nugget = 10). (B) High spatial autocorrelation (nugget = 0). Species exhibit a specialist–generalist trade-off mediated by the tolerance–fecundity trade-off, with fecundity drawn from $Y \sim \text{lognormal}(\mu = 0, \sigma_Y = 2.0)$ and species with lower fecundity tolerating a wider range of habitats. Note that interspecific variation in $Y$ is substantially larger than the main text baseline to better reflect seed production variation. Unless otherwise noted, parameters are $a = 0.5$, $\nu = 0.3 \times 10^{-5}$, and habitat range = 0.5.

##### 3.2 Simulations and results

I ran two sets of simulations. First, I conducted simulations analogous to those examined in Fig. 3 of the main text (restricted here to the global dispersal case for simplicity). Instead of varying $\sigma_h$ between runs, I varied the steepness parameter $Z$. In these simulations, I set $b_0 = 0.05$ and explored $Z \in \{0, 15, 25, 50, 150\}$, spanning cases from no habitat filtering to nearly step-like tolerance limits. Fecundities $Y_i$ were drawn from a lognormal distribution and then sorted in decreasing order to assign ranks ($R_i = 1$ most fecund), thereby implementing the fecundity–tolerance trade-off. Note that I set $Y \sim \text{lognormal}(\mu = 0, \sigma_Y = 2.0)$ rather than $\sigma_Y = 0.3$ (as in the main text), based on evidence that interspecific variation in seed fecundity *per se* is of this order (Chisholm & Fung, 2020).

Species richness increased with both $Z$ and $M$ (the JC-effect scale), consistent with the patterns shown in the main text. As before, species richness was substantially greater when habi-

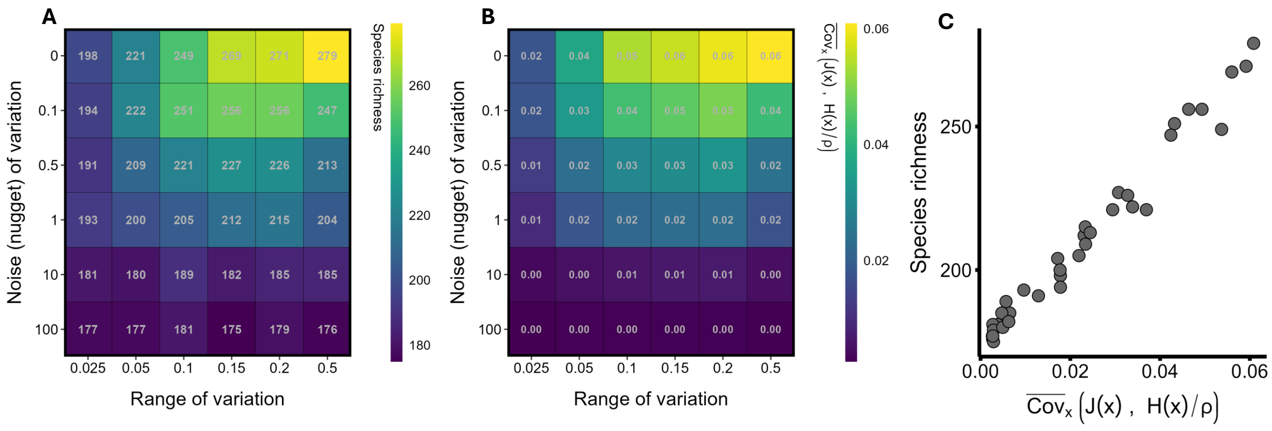

Figure 3: **When both JC-effects and habitat partitioning operate, species richness increases with abiotic habitat spatial autocorrelation, as reflected by the JC–HP covariance (under the tolerance-fecundity trade-off).** All panels depict simulations in which the range (x-axis) and noise (nugget, y-axis) of habitat variation were varied. (A) shows how range and nugget influence species richness. (B) shows how they influence the JC–HP covariance. (C) shows the relationship between species richness and the JC–HP covariance across simulations. Species exhibit a specialist–generalist trade-off mediated by the tolerance–fecundity trade-off. Here, $Y \sim \text{lognormal}(\mu = 0, \sigma_Y = 2.0)$ (large interspecific variation in fecundity), and species exhibit Z-shaped habitat responses in which seedling survival declines rapidly past a threshold habitat type. Species with lower fecundity ($Y$) tolerate a wider range of habitats (see Appendix for mathematical details). Unless otherwise noted, parameters are $a = 0.5$, $\nu = 0.3 \times 10^{-5}$, $Z = 150$, $M = 7$, and dispersal is global.

tats were highly spatially autocorrelated compared to when they exhibited low autocorrelation (Fig. 2).

Second, I ran simulations analogous to those in Fig. 4 of the main text, which examine how spatial autocorrelation influences species richness and the JC–HP covariance. As in the main text, both species richness and the JC–HP covariance increased with the degree of spatial autocorrelation (Fig. 3).

#### 4 Interspecific variation in baseline JC-effect strength

In the main text, I note that results are qualitatively unchanged when species differ in their baseline JC-effect strength $a$. To demonstrate this, I ran simulations in which $a$ varied between species $i = 1, 2, .., N$:

$$a_i \sim \text{Gamma}\Big(k = \tfrac{1}{\text{CV}^2},\ \theta = \tfrac{\bar{a}}{k}\Big), \tag{102}$$

with target mean $\bar{a} = 0.5$ and coefficient of variation CV $= 0.5$. This parameterization gives $k = 4$ and $\theta = 0.125$, so species-specific JC strengths $a_i$ were drawn from Gamma$(4, 0.125)$ across $S = 500$ species. In practice, this distribution produced values in the range $\sim 0.05 \leq a \leq 2$.

All other simulation parameters matched those in Fig. 4 of the main text (grid size = 175,

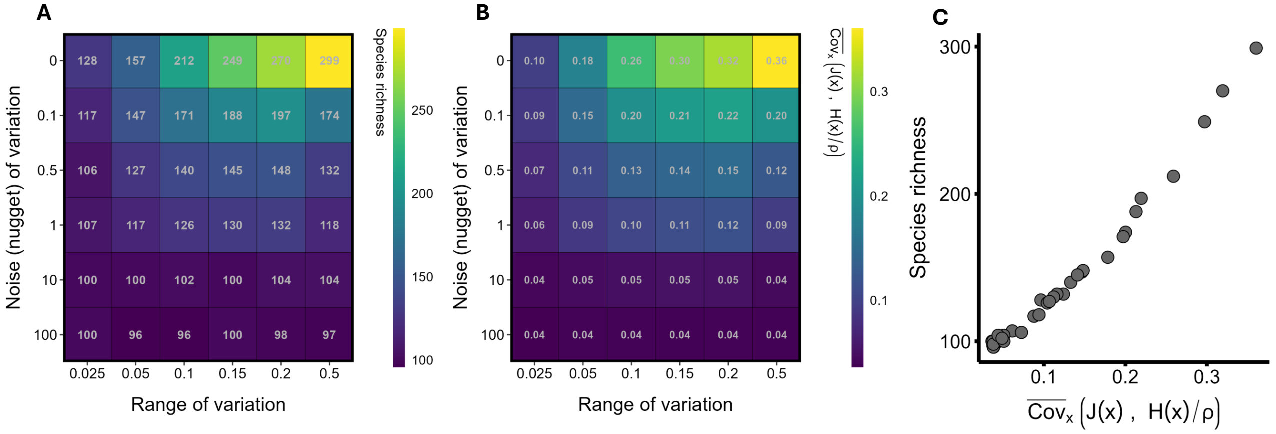

Figure 4: **When both JC-effects and habitat partitioning operate, species richness increases with abiotic habitat spatial autocorrelation, as reflected by the JC–HP covariance (when there is inter-specific variation in $a$).** All panels depict simulations in which the range (x-axis) and noise (nugget, y-axis) of habitat variation were varied. (A) shows how range and nugget influence species richness. (B) shows how they influence the JC–HP covariance. (C) shows the relationship between species richness and the JC–HP covariance across simulations. Species exhibit interspecific variation in JC-effect strength $a$, drawn from a gamma distribution with mean $\overline{a} = 0.5$ and coefficient of variation $= 0.5$ (yielding $\sim 0.05 \leq a \leq 2$). Unless otherwise noted, parameters are $\nu = 0.3 \times 10^{-5}$, $\sigma_h = 0.02$, $Y \sim \text{lognormal}(\mu = 0, \sigma_Y = 0.3)$, $M = 7$, and dispersal is global.

$n_{\text{generations}} = 10{,}000$, $\nu = 0.3 \times 10^{-5}$, $\sigma_h = 0.02$, $M = 7$, global dispersal), with spatial autocorrelation controlled by $\lambda$ and the range of habitat variation as described in the main figure.

Introducing interspecific variation in $a$ reduced equilibrium species richness relative to the constant-$a$ case. Nevertheless, richness remained high, and—most importantly—the qualitative patterns were the same as those shown in the main text (Fig. 4).

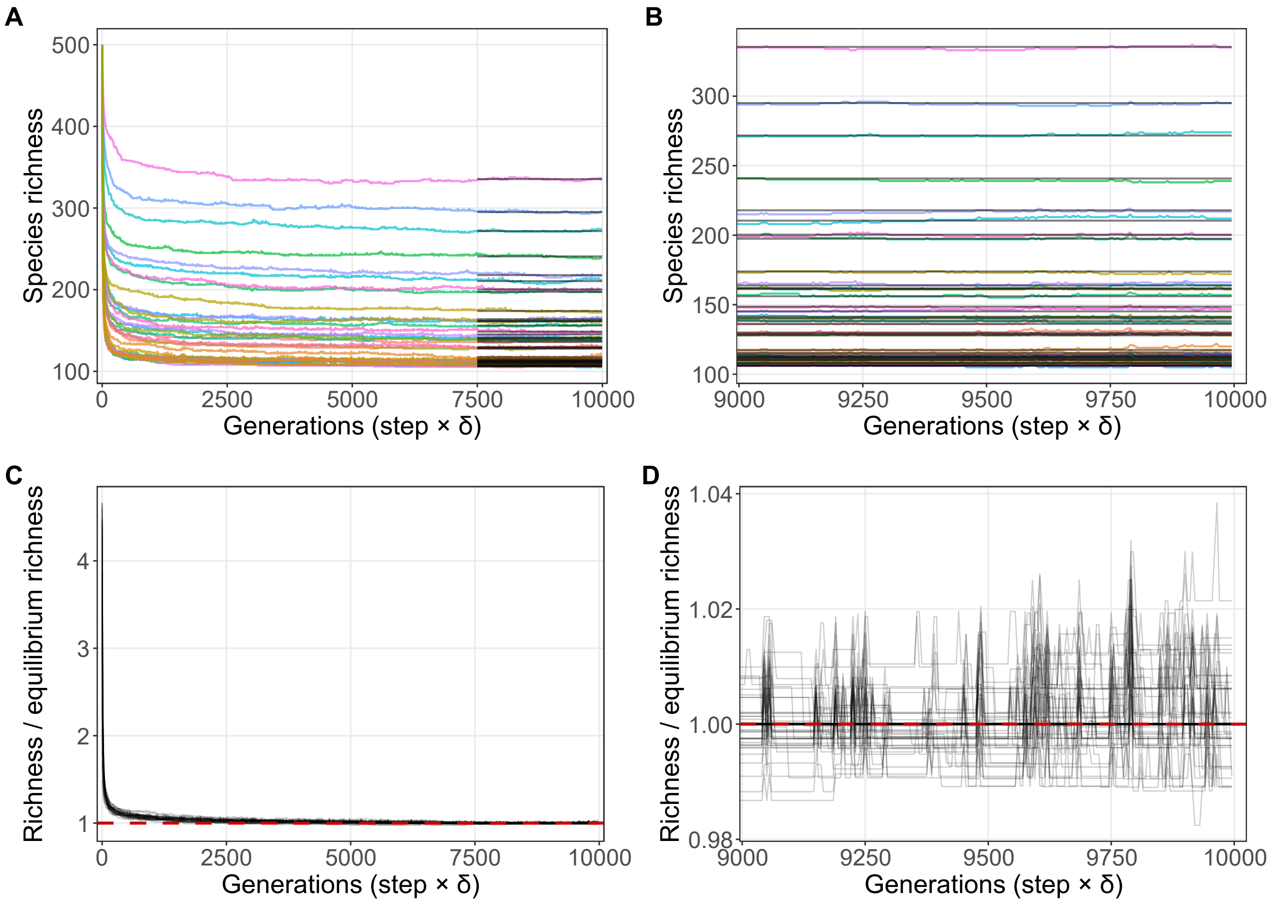

Figure 5: **Richness trajectories indicate dynamic equilibrium for the simulations shown in Figure 3.** (A) Species richness through time (generations; time steps $\times\,\delta$) for each simulation (colored lines). Horizontal segments show the simulation-specific equilibrium richness, defined as the mean richness over the final $n = 500$ recorded output times. (B) Same as (A), zoomed to the final 1000 generations. (C) Normalized richness trajectories, obtained by dividing richness by the simulation-specific equilibrium richness; the dashed line at 1 indicates the equilibrium value under this normalization. (D) Same as (C), zoomed to the final 1000 generations. Note that the balance between extinction and immigration over the last 1000 generations.

#### 5 Confirmation of dynamic equilibrium

To confirm that simulations approach a stationary richness regime within the simulated time horizon, I tracked species richness through time, defining richness at each recorded output time as the number of species with non-zero occupancy (i.e., occupying at least one grid cell). Time is reported in generations (time steps $\times\,\delta$). Supplementary Fig. 5 shows this diagnostic for the simulations depicted in Fig. 3 of the main text; the equilibration patterns shown here are typical of those observed across simulation experiments. (Simulation state was recorded every 25 time steps.)

In Supplementary Fig. 5A–B, each line shows richness through time for a single simulation (y-axis: species richness; x-axis: generations). For each simulation, the horizontal segment indicates the simulation-specific equilibrium richness, defined as the mean richness over the final $n = 500$ recorded output times. Panel 5B zooms to the final 1000 generations to emphasize late-time stationarity.

In Supplementary Figure 5C–D, I plot normalized trajectories obtained by dividing richness at each time point by the simulation-specific equilibrium richness (estimated from the same final $n = 500$ recorded output times). The dashed horizontal line at 1 marks the equilibrium value under this normalization, and Panel D zooms to the final 1000 generations. Together, these diagnostics show that richness approaches and remains close to its terminal level over the latter portion of the run.

#### 6 Analysis of approximations

Using the simulation outputs shown in Figs. 3A, 3B, and 4, I evaluated the accuracy of the analytical approximation for the invader invasion criterion by comparing the simulation-based value $\tilde{\lambda}_{i,\text{exact}}$ to the corresponding estimate $\tilde{\lambda}_{i,\text{approx}}$ obtained by evaluating Eq. 47 (equivalently, Eq. (6) in the main text) with inputs measured from each simulated community.

For each parameter combination and stochastic realization (i.e., each community simulation run), I simulated the resident community to its long-run state and recorded the final spatial configuration of resident adults. Holding the final resident community fixed, I computed $\tilde{\lambda}_{i,\text{exact}}$ by numerically evaluating Eq. 19 for each invasion attempt: I drew an invader's traits (including $Y_i$ and $h_{\text{opt},i}$), computed the resulting spatial fields in the numerator, and evaluated the denominator using the resident species' contributions from the realized end-of-simulation adult configuration. I then averaged the bracketed quantity over space (i.e., over grid cells) and subtracted 1. In parallel, for the same invader and resident configuration, I computed $\tilde{\lambda}_{i,\text{approx}}$ by substituting numerically measured quantities from that resident configuration into Eq. 47. This procedure treats each community simulation run as a single resident-community replicate, with many invasion attempts nested within it, and it implicitly assumes the resident community is at (or near) dynamic equilibrium.

I show: (i) an invader-level visualization for each replicate (the set of $\tilde{\lambda}_i$ values produced per replicate) (Fig. 6); (ii) replicate-level mean comparisons $\overline{\tilde{\lambda}}_{\text{exact}}$ vs. $\overline{\tilde{\lambda}}_{\text{estimate}}$ (Fig. 7A); (ii) approximations of $\overline{h(x)}$, Eq. 78 contrasting with Eq. 86 while holding other numerically measured inputs fixed (Fig. 7B-C); and (iii) comparisons of simulation-measured $\mathbb{E}[J(x)]$ to the two approximations used in the main text (randomly distributed adults, Eq. 7; moments approximation, Eq. 8) (Figs. 7D-E).

Overall, $\tilde{\lambda}_{i,\text{estimate}}$ closely matches $\tilde{\lambda}_{i,\text{exact}}$ across replicates, with the key agreement occurring near the invasion threshold: on average $\tilde{\lambda}_{i,\text{estimate}} \approx \tilde{\lambda}_{i,\text{exact}}$ when $\tilde{\lambda}_{i,\text{exact}} \approx 0$ (Fig. 6; Fig. 7A), despite appreciable noise at the level of individual invasion attempts, resulting in few false positives or false negative predictions. For $\overline{h(x)}$, both Eq. 86 and Eq. 78 are quite accurate, and perform similarly (Fig. 6C vs. Fig. 7B); the latter, however, shows notable outliers in low-richness replicates (note the outliers in the upper-left area in B). Finally, both approximations for $\mathbb{E}[J(x)]$ (Eqs. 7–8, main text) are supported over the parameter ranges explored (Figs. Fig. 7D-E). However, assuming species to be randomly distributed in space (Eq. 7, main text) tends to overestimate $\mathbb{E}[J(x)]$ because (at least, in many simulations in Figs. 3A-3B, and 4), species are aggregated in space.

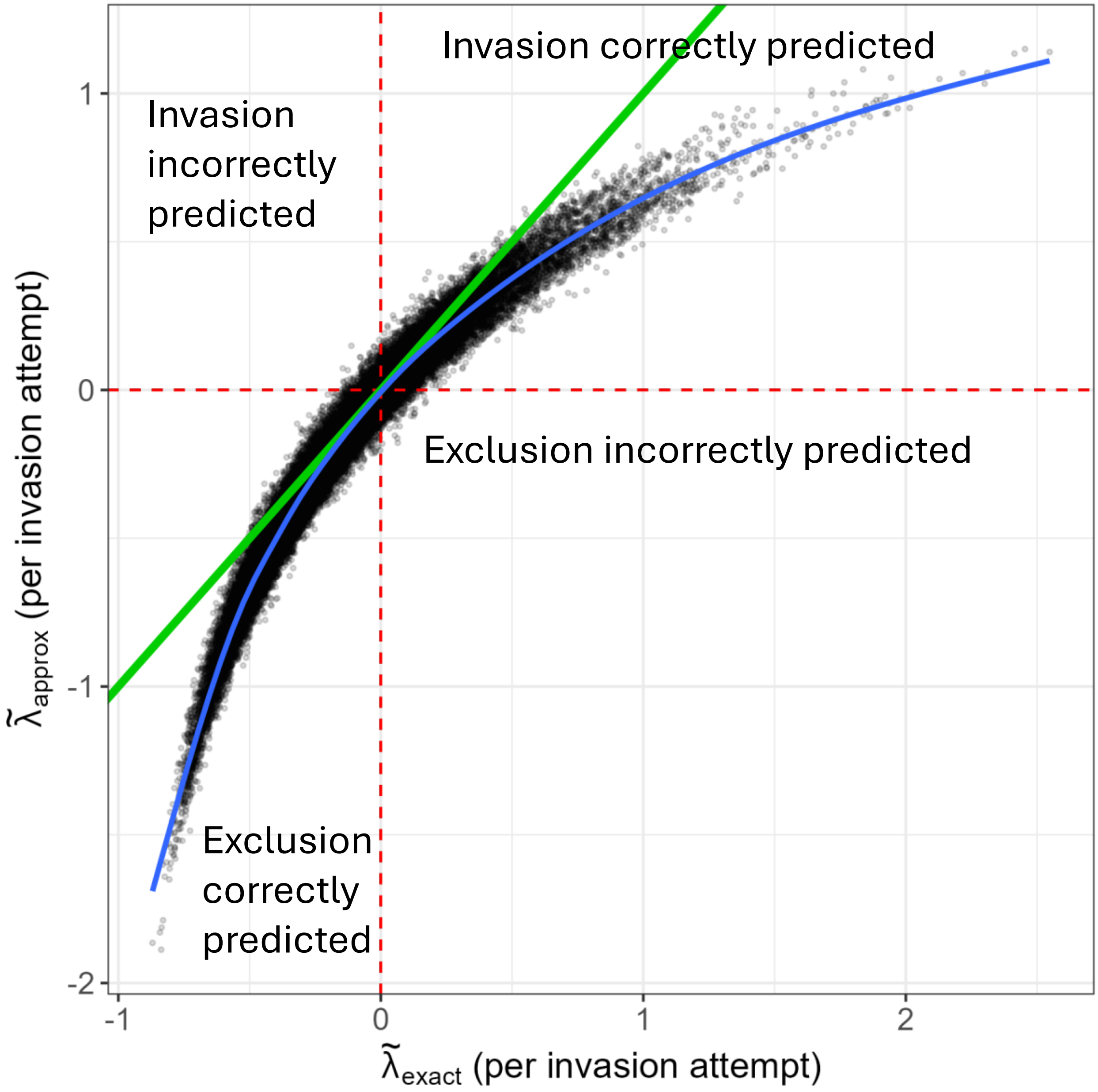

Figure 6: **Approximate and exact invasion criteria are closely aligned throughout invasion attempts.** The $x$-axis shows the simulation-based exact criterion, $\tilde{\lambda}_{i,\mathrm{exact}}$, and the $y$-axis shows the corresponding approximation, $\tilde{\lambda}_{i,\mathrm{approx}}$ (Eq. 47). Points show invasion attempts for invaders with randomly drawn fitness $Y_i$ and habitat optimum $h_{\mathrm{opt},i}$; for each simulated resident community, 500 invasion attempts were evaluated. Simulations are from Figs. 3A, 3B, and 4 presented in the main text. Red dashed lines mark the invasion boundaries $\tilde{\lambda}_{i,\mathrm{approx}} = 0$ (horizontal) and $\tilde{\lambda}_{i,\mathrm{exact}} = 0$ (vertical). The green line shows the 1:1 relationship, and the blue curve shows a spline fit summarizing the mean trend.

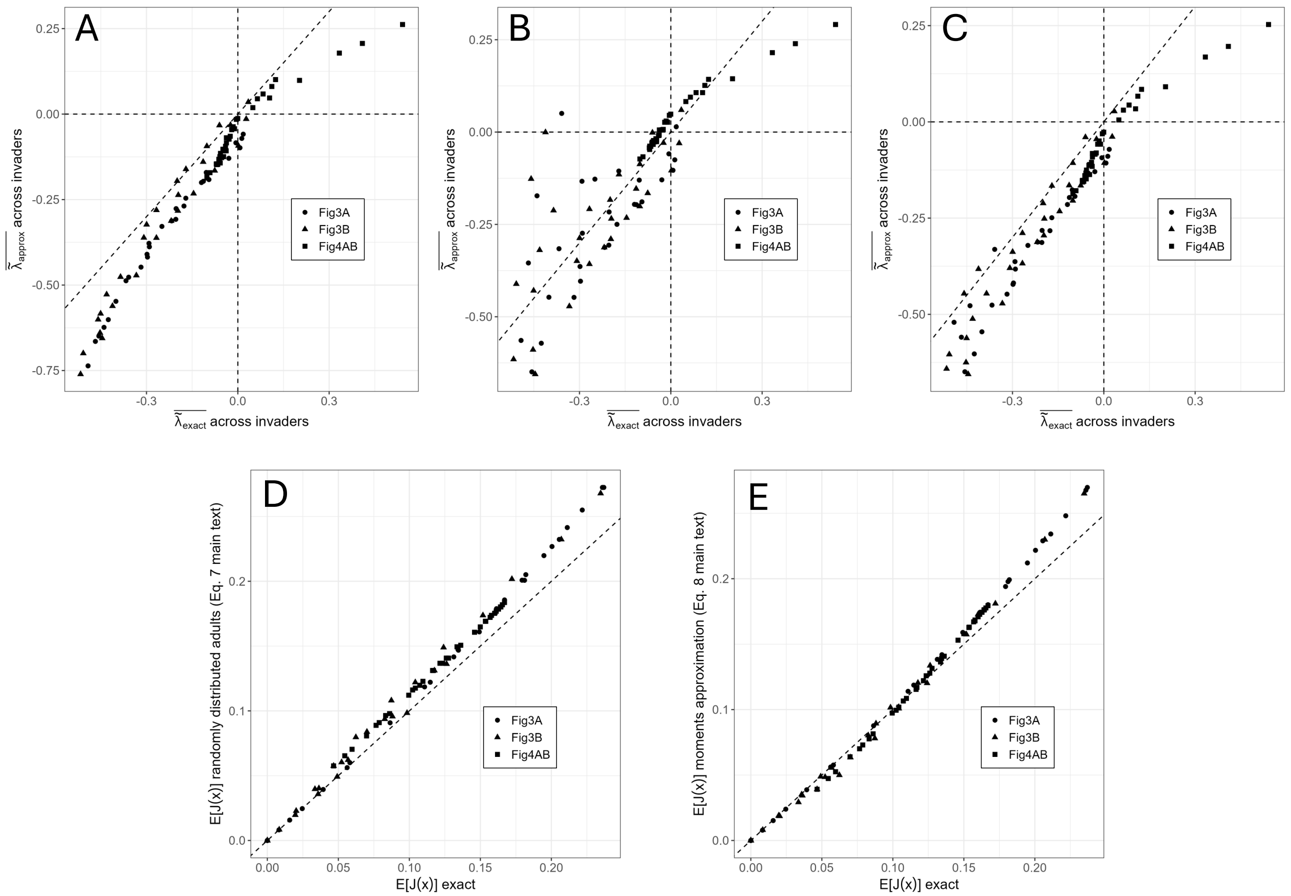

Figure 7: **Across simulation sets, alternative approximations typically yield similar comparisons to exact values.** Points denote simulation replicates from the runs shown in Figs. 3A, 3B, and 4 (circles, triangles, squares, respectively). Panels A–C compare replicate-level means across 500 invasion attempts (random draws of $Y_i$ and $h_{\text{opt},i}$ per replicate): the $x$-axis is $\bar{\bar{\lambda}}_{i,\text{exact}}$ across invaders and the $y$-axis is $\bar{\bar{\lambda}}_{i,\text{approx}}$ across invaders (Eq. 47). (A) uses numerically evaluated inputs from the realized end-of-simulation adult spatial configuration, including $\overline{\mathbb{E}_x[J(x)]}$ and $\overline{\text{Cov}_x}\Big(J(x), \frac{H(x)}{\mu_{H(x)}}\Big)$, and computes $\overline{h(x)}$ directly from Eq. 42. (B) is identical to (A) except $\overline{h(x)}$ is approximated using Eq. 78 (notable outliers occur primarily in low-richness replicates where this approximation is least accurate). (C) is identical to (A) except $\overline{h(x)}$ is approximated using Eq. 86. Panels D–E validate approximations for $\overline{\mathbb{E}_x[J(x)]}$: the $x$-axis shows $\overline{\mathbb{E}_x[J(x)]}$ computed numerically from the final adult spatial distribution, while the $y$-axis shows (D) the randomly distributed adults approximation (Eq. 7, main text) and (E) the moments approximation (Eq. 8, main text). Dashed diagonal lines indicate the 1:1 relationship (upon which the approximation is identical to the exact quantity); dashed vertical and horizontal lines in A–C mark $\bar{\bar{\lambda}}_{i,\text{exact}} = 0$ and $\bar{\bar{\lambda}}_{i,\text{approx}} = 0$, respectively.

#### 7 Parameter tables

Table 1: Notation and parameters used in the **main text**.

| Symbol / term | Meaning | Notes / domain |
|---|---|---|
| *Indices, space, and community sizes* | | |
| $i, j, k$ | Species indices (with $i$ often the focal/rare invader). | $1, \ldots, N$ |
| $t$ | Discrete time step. | |
| $x$ | Patch (site) index / spatial location. | |
| $P$ | Number of patches in the local landscape. | e.g. $X \times X$ |
| $N$ | Number of species currently present locally. | |
| $L$ | Number of species in the regional species pool. | |
| $p_i(t)$ | Adult frequency of species $i$ (fraction of patches occupied). | $\sum_i p_i(t) = 1$ |
| *Demography, dispersal, and recruitment* | | |
| $\delta$ | Adult mortality probability per time step (fraction of patches vacated and recolonized). | $\delta = [0, 1]$ |
| $\nu$ | Immigration probability (colonist drawn from regional pool rather than local propagule pool). | $\nu = 0.3 \times 10^{-5}$ |
| $Y_i$ | Intrinsic fitness of species $i$ (multiplicative composite; e.g. fecundity $\times$ baseline offspring survival $\times$ low-density recruitment). | $Y_i > 0$ |
| $\sigma_Y$ | Log-scale standard deviation for intrinsic fitness variation across species. | lognormal draw |
| $d_i(x)$ | Dispersal pressure of species $i$ arriving at patch $x$ (kernel-weighted adult contributions). | relative supply |
| $\sigma_D$ | Dispersal scale parameter (kernel width). | shared in sims |
| $S_i(x)$ | Surviving propagules of species $i$ at patch $x$ after dispersal and juvenile survival filters. | enters lottery |
| *Habitat filtering and landscape generation* | | |
| $E(x)$ | Habitat value at patch $x$. | rescaled to $[0, 1]$ |
| $h_{\text{opt},i}$ | Habitat optimum (preferred habitat) of species $i$. | $h_{\text{opt},i} \in [0, 1]$ |
| $\sigma_h$ | Niche breadth (width of habitat response). | smaller = narrower |
| $H_i(x)$ | Habitat performance / survival multiplier of species $i$ at patch $x$. | Gaussian response |
| sill | Total variance of the Gaussian random field at large distances. | fixed at 1 |
| range | Distance where sites become effectively uncorrelated (variogram reaches 95% of sill). | scaled to landscape |
| nugget | Microscale white-noise variance added per patch (local roughness). | |
| *Janzen–Connell (JC) effects* | | |
| $M$ | Linear size of Moore neighborhood of influence. | neighborhood $M \times M$ |
| $A_i(M(x))$ | Number of conspecific adults of species $i$ within the $M \times M$ neighborhood around $x$. | neighborhood count |
| $a_i$ (or $a$) | Per-conspecific JC-effect strength (cumulative hazard per conspecific adult). | $a_i > 0$ |
| $J_i(x)$ | JC-effect mortality probability at patch $x$. | e.g. $1 - e^{-a_i A_i(M(x))}$ |
| *Analytical invasion approximation (main-text expressions)* | | |

*(continued)*

| Symbol / term | Meaning | Notes / domain |
|---|---|---|
| $\lambda_i$ | Finite rate of increase of rare invader $i$ (ratio of frequencies across steps). | $p_i(t+1)/p_i(t)$ |
| $\tilde{\lambda}_i$ | Weighted growth rate used for invasion criterion. | $(\lambda_i - 1)/\delta$ |
| $\overline{\ln Y}$ | Resident-mean log intrinsic fitness. | mean over residents |
| $\overline{\mathbb{E}_x[J(x)]}$ | Resident-mean spatial mean JC mortality probability. | mean over spp. and space |
| $\overline{h(x)}$ | Habitat term in the invasion decomposition. | main text def. |
| $\mu_{H(x)}$ | Spatial mean habitat response. | $\mathbb{E}_x[H(x)]$ |
| $\sigma^2_{H(x)}$ | Spatial variance of habitat responses across patches. | |
| $\mathrm{CV}^2_{H(x)}$ | Squared coefficient of variation of habitat response. | $\sigma^2_{H(x)}/\mu^2_{H(x)}$ |
| $\bar{\rho}$ | Average correlation of habitat responses across species. | enters $\overline{h(x)}$ |
| $\sigma_{J(x)}$ | Spatial standard deviation of JC mortality probability. | used in bound |
| $\overline{\mathrm{Cov}_x}\Big(J(x),\, \frac{H(x)}{\mu_{H(x)}}\Big)$ | Community-mean spatial covariance between JC pressure and (normalized) habitat suitability. | "JC–HP covariance" |
| $\kappa_{i,M}$ | Scale-$M$ aggregation index (variance-to-mean ratio of conspecific neighborhood counts). | $\kappa > 1$ aggregated |
| $\overline{\kappa}_M$ | Community mean of $\kappa_{i,M}$ at scale $M$. | |
| $\mu_{i,M}$ | Mean conspecific neighborhood count for species $i$ at scale $M$. | $\mathbb{E}[A_i(M(x))]$ |
| *Aggregation metric used for simulation summaries* | | |
| $\psi$ | Point-process intensity (adult density; expected adults per unit area). | |
| $K(r)$ | Ripley's $K$-function (expected neighbors within $r$ is $\lambda K(r)$). | CSR: $K(r) = \pi r^2$ |
| $g(r)$ | Pair-correlation function. | $g(r) = K'(r)/(2\pi r)$ |
| $r$ | Distance scale at which spatial clustering is evaluated. | |

Table 2: Additional notation used **only in the Supplementary Information**.

| Symbol / term | Meaning | Notes / domain |
|---|---|---|
| *Landscape geometry and dispersal-kernel implementation* | | |
| $X$ | Linear grid dimension, so $P = X \times X$. | integer |
| $y$ | Source patch index (used in dispersal sums into focal patch $x$). | |
| $s(y)$ | Species identity occupying patch $y$. | |
| $\mathbf{1}\{s(y) = i\}$ | Indicator: 1 if patch $y$ is occupied by species $i$, else 0. | |
| $\mathrm{dist}(x, y)$ | Chebyshev distance between patches $x$ and $y$. | integer shells |
| $z$ | Chebyshev distance shell index. | $z \in \{0, 1, \dots\}$ |
| $\zeta$ | Euclidean distance (used in the underlying isotropic kernel). | $\zeta \geq 0$ |
| $k(\zeta; \sigma_D)$ | Radial dispersal kernel density (continuous form used to build discrete kernel). | e.g., Gaussian/Rayleigh form |
| $E(z)$ | Discrete dispersal probability assigned to shell $z$ (per destination cell). | derived from $k$ |
| $z_{\max}$ | Maximum shell handled explicitly when discretizing the kernel. | integer |
| $D_{\mathrm{Lim}}$ | Total kernel mass captured by explicit shells $\{0, \dots, z_{\max}\}$. | $0 \leq D_{\mathrm{Lim}} \leq 1$ |
| $1 - D_{\mathrm{Lim}}$ | Remaining tail mass redistributed uniformly across the landscape. | long-distance tail |
| *Vulnerable-period survival bookkeeping (continuous-time within a step)* | | |
| $f_i$ | Fecundity of species $i$ (propagules produced per adult per step). | $f_i > 0$ |
| $\tau$ | Duration of the vulnerable period (juvenile stage over which mortality acts before recruitment). | time units of model |
| $m_i$ | Baseline density-independent mortality hazard accumulated over the vulnerable period. | $m_i \geq 0$ |
| $\ell_i(x)$ | Density-independent mortality rate at patch $x$ (baseline + habitat component). | rate |
| $j_i(x, t)$ | Density-dependent (JC) mortality rate experienced at patch $x$ during the vulnerable period. | rate |
| $S_i(x, t)$ | Surviving propagules at patch $x$ as a function of within-step time $t \in [0, \tau]$. | SI derivations |
| $S_i(x, 0)$ | Initial propagule number entering the vulnerable period. | typically $d_i(x) f_i$ |
| *Neighborhood/set notation used in derivations* | | |
| $\mathcal{N}_M(x)$ | Set of patches in the $M \times M$ Moore neighborhood around $x$. | includes focal window |
| $A_j(x)$ | Conspecific neighborhood count for species $j$ around $x$. | random variable |
| $\sigma^2_{j,M}$ | Variance of $A_j(x)$ at neighborhood scale $M$. | |
| *Dispersal-limitation approximation used in SI* | | |
| $J_i^L(x)$ | Additional survival reduction for propagules that remain near the parent (local-parent JC effect). | SI-only notation |
| $d$ | Fraction of propagules that escape local-parent effects in the simplified approximation. | $0 \leq d \leq 1$ |
| *Moment-expansion / covariance-structure notation* | | |
| $C_{jk}$ | Spatial covariance between habitat responses $H_j(x)$ and $H_k(x)$ (or related cross-moments, as defined in SI). | SI derivations |

*(continued)*

| Symbol / term | Meaning | Notes / domain |
|---|---|---|
| $\mu_{j,M}$ | Mean of $A_j(x)$ (conspecific neighborhood count) for species $j$ at scale $M$. | |
| $\mathrm{Corr}_x(\cdot,\cdot)$ | Spatial correlation across patches $x$. | SI derivations |
| *Parameter heterogeneity specifications (SI-only)* | | |
| $\overline{a}$ | Target mean JC-effect strength across species when drawing $a_i$. | $> 0$ |
| CV | Coefficient of variation used to control heterogeneity in $a_i$ draws. | $> 0$ |
| $k$ | Gamma shape parameter for $a_i$ distribution. | $k = 1/\mathrm{CV}^2$ (SI) |
| $\theta$ | Gamma scale parameter for $a_i$ distribution. | $\theta = \overline{a}/k$ (SI) |
| *Alternative habitat-tolerance specification in SI ("Model specifications" section)* | | |
| $S$ | Number of species in that SI-only ranked-tradeoff specification. | distinct from $S_i(x)$ |
| $R_i$ | Fecundity rank of species $i$ (used to impose a fecundity–tolerance tradeoff). | $1, \ldots, S$ |
| $b_0$ | Baseline tolerance intercept in the SI logistic habitat establishment function. | $b_0 \in [0, 1]$ (SI) |
| $Z$ | Steepness parameter controlling how sharply establishment probability changes along the habitat gradient. | $Z > 0$ |
| $L_i$ | SI-only tolerance/limit quantity appearing in the ranked-tradeoff construction (as defined there). | see SI definition |
| *Simulation bookkeeping (SI-only)* | | |
| $n_{\mathrm{generations}}$ | Number of generations/time steps simulated. | integer |
